# Identification of prognostic biomarkers for suppressing tumorigenesis and metastasis of Hepatocellular carcinoma through transcriptome analysis

**DOI:** 10.1101/2022.07.20.500640

**Authors:** Divya Mishra, Ashish Mishra, M.P. Singh

## Abstract

Cancer is one of the deadliest diseases developed through tumorigenesis and could be fatal if reached at metastatic phase. The novality of present investigation is to explore the prognostic biomarkers in hepatocellular carcinoma (HCC) that could develop glioblastoma multiforme (GBM) due to metastasis. The analysis was done using RNA-Seq datasets for both HCC (PRJNA494560 and PRJNA347513) and GBM (PRJNA494560 and PRJNA414787) from Gene Expression Omnibus (GEO). This study identified 13 hub genes found to be overexpressed in both GBM and HCC. Promoter methylation study showed these genes to be hypomethylated. Validation through genetic alteration and missense mutations resulted in chromosomal instability leading to improper chromosome segregation causing aneuploidy. A 13-gene predictive model was obtained and validated using KM plot. These hub genes could be prognostic biomarkers and potential therapeutic targets, inhibition of which could suppress tumorigenesis and metastasis.

## Introduction

Cancer is a complex disease caused due to uncontrolled division and growth of cells, categorized according to the progression in organs such as breast cancer, blood cancer, colon cancer, liver cancer, etc. Liver cancer, also known as hepatocellular carcinoma (HCC), has nowadays become a common cancer type, with approximately 830000 deaths in 2020 alone (Gao et al., 2021). Tumorigenesis or transformation of normal cells into cancerous cells often results in uncontrolled cell proliferation, metastasis, and apoptosis evasion (Cao, 2017). Metastasis of cancer cells occurs through blood vessels, and lymph nodes and account for the development of other types of cancers (Fares et al., 2020). Its occurrence is common in hepatocellular carcinoma (HCC) patients’ undergone surgery (Zhu et al., 2018). Most of the patients’ are diagnosed during late phase of the disease. Significant advancements in early disease diagnosis through standard interventions such as radiation, surgery, personalized strategies, chemotherapy have been developed in the past few decades. The aggregative five-year survival rate of HCC and GBM remains destructive due to their molecular heterogeneity and invasive behavior. It has been found in some studies that brain metastases from HCC are less frequent (0.2%-2.2%) and resulted in poorer survival of patients (Jiang et al., 2012). Even though this rate is less, but still it is certain that 10%-45% of different cancer types from liver, lung and other body parts metastasize to brain (Lah et al., 2019). This process of metastasis can be dealt with using biomarkers. Biomarkers are biological molecules found in body fluids, blood samples, or tissues and these signify a particular condition or disease. In different cancer types, metastatic biomarkers helps in detecting the early stages of tumor spread, recurrence probability, and also in predicting the favorable sites of metastasis (Brinton et al., 2012). Once metastatic cancer is detected, further we need to identify the DNA or RNA-based biomarkers that could allow for personalized therapy resulting in significantly positive survival outcomes in patients (Dawood, 2010). Some biomarkers as CDK4, PTEN, and ERBB2 were found as potential indicators for diagnosis and prediction ofmetastatic breast cancer (Fasching et al., 2015). Likewise, overexpression of EGFR in metastatic non-small cell lung cancer (NSCLC) makes it a prognostic biomarker (Scott et al., 2008). In the past few years, the fast growth of *in silico* approaches such as next-generation sequencing has enabled insight into carcinogenesis and progression of distinct cancer (Park et al., 2021). High-throughput platforms have been extensively used in prognosis prediction, histological identification, early diagnosis, disease resistance analysis and molecular classification. Long noncoding RNAs (lncRNAs), microRNAs (miRNAs), differentially expressed genes (DEGs) and differentially methylated CpG sites can potentially serve as valuable HCC biomarkers (Tang et al., 2020). Few oncogenic lncRNAs such as LASP1-AS, MALAT1, HOTAIR, and NORAD acted as potential biomarkers in case of HCC. Similarly, some tumor-suppressor genes viz. DGCR5, MIR22HG, and HOTAIRM1 also found as potential biomarkers for HCC (Yuan et al., 2021). An example of miRNAs includes miR-125a-5p that was upregulated in patients having HCV-associated hepatocellular carcinoma (Oura et al., 2019). Similarly, the overexpression of some differentially expressed genes such as CDC20, BUB1B, AURKA, CCNA2, BUB1 were found responsible for poor progression and high mortality in patients suffering from HCC (Li et al., 2021).The transcriptome analysis has disclosed the cancer molecular mechanisms. Meanwhile, few reports have been introduced to identify the candidate biomarkers related to HBV-HCC with combined datasets (Tang et al., 2020). In this study, the transcriptome analysis was carried out using RNA-Seq datasets to identify the differentially expressed genes (DEGs) that play a vital role as potential prognostic biomarkers in case of metastatic HCC and GBM. One of the most important biological processes obtained from DAVID database using DEGs, chromosome segregation, has a prominent role in tumorigenesis as any chromosomal instability causes genetic instability due to dysregulated chromosome segregation (Zhang et al., 2018). Similarly, cell cycle, which is an important KEGG pathway, was obtained. Any aberrant change in this cycle may also result in tumorigenesis. Hence, its regulators could be treated as potential anticancer therapeutic targets (Liu et al., 2021).

Another predominant feature related to tumor development and progression is alteration in DNA methylation. DNA hypomethylation is more prominent with tumorigenesis or malignancy than hypermethylation (Ehrlich et al., 2006; Qu et al., 1999; Roman-Gomez et al., 2008; Park et al., 2009). It has been found that genomic instability occurs due to DNA hypomethylation in case of HCC (Tischoff et al., 2008). This instability causes activation of oncogenes such as antigen family A1 (*MAGEA1*) (Chakravarthi et al., 2016). Genetic alterations in the form of mutations and DNA copy number alterations (CNAs) were also identified as critical features of HCC tumorigenesis and metastasis. A study found that missense mutation in NUF2 gene was linked to cancer development, and hence, its inhibition resulted in suppression of tumor growth leading to cancer cells apoptosis (Kamburov et al., 2015). Copy number alterations are present in 90% of solid tumors and play a prominentrole in activating oncogenes and inactivating tumor suppressor genes by altering the dosage and structure ofgenes (Court et al., 2020). As CNAs outline pivotal genetic events that drive tumorigenesis, such genetic alterations have the potential as predictive factors (Bergamaschi et al., 2006). Post-translational modifications (PTM) viz. phosphorylation, acetylation, Ubiquitination, methylation, sumoylation, etc. also play vital role in tumorigenesis of different cancer types, particularly in breast cancer (Theivendran et al., 2020). Mutation in Aurora Kinase A (AURKA) in HCC through direct phosphorylation of Pkinase promoted tumorigenesis and subsequently metastasis (Ardito et al., 2017). Likewise, Ubiquitination, another PTM plays a vital role in administering the control of substrate degradation, which is required for the proper functioning of the cell cycle and any abberancy in this process will hamper normal cell functioning leading to cancer development and later probable metastasis (Deng et al., 2020). In this study, aberrant Ubiquitination in Pkinase led to mutation in AURKA gene, and this abnormal overexpression resulted in tumorigenesis and later stage metastasis of HCC. This study, therefore involved the identification of differentially expressed genes that were overexpressed in both GBM and HCC. The 13 hub genes obtained were further validated through promoter methylation, mutation and genetic alterations analysis proved their potential to be prognostic biomarkers. The survival analysis of all these hub genes showed poorer survival rates among metastatic HCC and GBM patients.

## Materials and Methods

The datasets for both GBM and HCC was taken from Gene Expression Omnibus (GEO). For GBM (normal samples-PRJNA494560 transcriptomic data with paired-end sequencing performed on Illumina HiSeq 3000 (Homo Sapiens) platform, and tumor samples-PRJNA347513, transcriptomic data with paired-end sequencing performed on Illumina HiSeq 2000 platform) and for HCC (normal samples-PRJNA494560 transcriptomic data with paired-end sequencing performed on Illumina HiSeq 3000 (Homo Sapiens) platform and tumor samples-PRJNA414787 transcriptomic data with paired-end sequencing performed on Illumina HiSeq 2000 (Homo Sapiens) platform) were taken. The method that was followed for carrying out this study is shown below in flowchart (see Figure 1).

**Figure 1.**
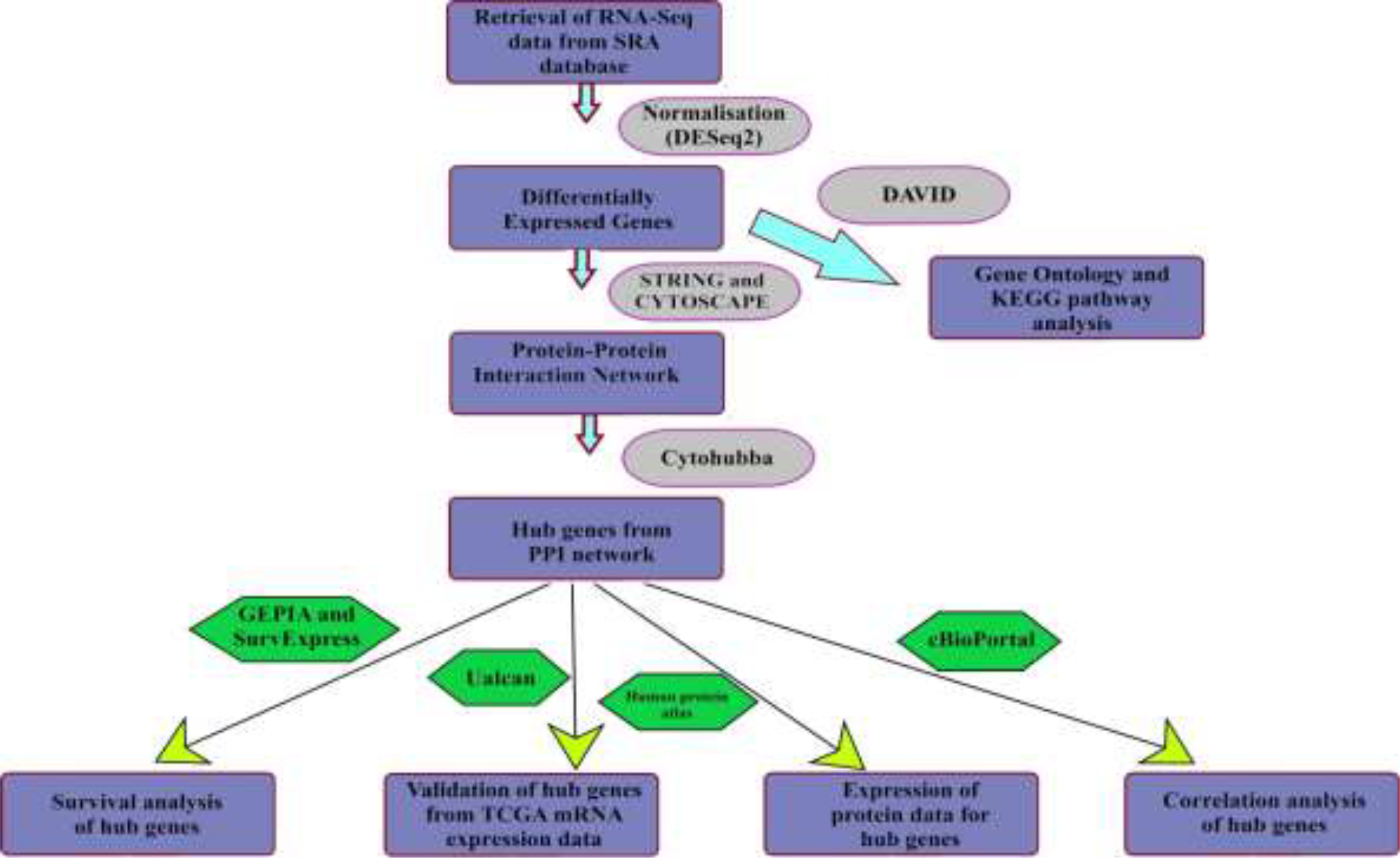
Flowchart showing the methodology followed in the present study

### Identification of common prognostic biomarkers in

#### HCC and GBMData Pre-processing

The data pre-processing of the raw reads was performed on Galaxy, an open source platform for analyzing genomic data (Afgan et al., 2018). Galaxy implements FastQC (Version 0.11.8), a quality control tool for high-throughput sequenced data, for conducting the quality assessment of raw reads and removing the adapter sequences, uncalled bases and low quality reads for improving the sequence quality through filtering and trimming. For this purpose, Cutadapt tool (v 3.2) is implemented. After obtaining the high-quality data from pre-processing, the next step that follows is alignment of reads against Human reference genome (GRch38/hg38). This is accomplished using STAR (Version 2.7.7a) which is an ultrafast universal RNA-seq alignment tool (Dobin et al., 2013). The mapped reads are subsequently quantified in a process called quantification, through featureCounts (subread Version 2.0.1) package (Liao et al., 2013). This step provides read counts per annotated gene. The normalized read count is further taken and eventually the statistical analysis is performed to obtain the differentially expressed genes (DEGs) between control and treated groups. It provides the quantitative changes in expression levels of genes. DESeq2 (Version 1.22.1) is a tool that performs this normalization process and is based on negative binomial distribution (Love et al., 2014). DEGs having FDR (p-value (adj)) <0.05 and |Log2FC| >2 are considered statistically significant.

### Protein-Protein Interaction Network Analysis and Identification of hub genes

The protein-protein interaction network is critical for understanding the cellular processes in diseased and normal states. This network provides the mathematical representations of the physical contacts between different proteins. This network was obtained by taking DEGs as input in the STRING database (Szclarczyk et al., 2020). The vertices constitute DEGs (proteins) and edges constitute the proteins interactions. The network was subsequently visualized through Cytoscape software (Shannon, P. et al., 2003). The confidence score was taken <0.4. The PPI enrichment value < 1.0e-16 indicated that the network has significant interactions. The modular analysis was obtained by implementing MCODE (Molecular Complex detection) plug-in of cytoscape. The parameters included degree cut-off =0.2, node score cut-off = 0.2, k-core = 2 and maximum depth = 100. The hub genes are then identified from the obtained module using Cytohubba plug-in. Hub genes are hugely interconnected genes and plays a critical role in PPI network. For this purpose, 5 different topologies i.e. Maximal Clique Centrality (MCC), Degree, Edge Percolated Component (EPC), Maximum Neighborhood Component (MNC), and Radiality were employed. The topmost 15 were considered as hub genes in all the 5 algorithms and the common 13 hub genes were then taken from these 5 topologies through venn diagram obtained from jvenn (Bardou et al., 2014).

### GO Component and Pathway Enrichment Analysis

Both GO and KEGG pathway enrichment analysis was obtained by providing common DEGs between GBM and HCC as input to DAVID database which is an online tool for functional enrichment analysis (Dennis Jr et al., 2003). For both GO term and KEGG pathway, the EASE value (modified Fisher Exact P-value), employed for measuring the gene-enrichment in annotation terms, was set to 0.1, and the count threshold to 2 (default value). The lesser this P-value is, the more enriched the GO terms or KEGG pathways are. The cut-off value for any term or pathway to be significant was set at p<0.05. REVIGO (Supek et al., 2011) was used subsequently for constructing the treemap for biological processes by entering GO ids of all the terms along with their respective p-values.

### Epigenetic Regulation of Gene Expression of Hub Genes by Promoter Methylation

DNA methylation is an epigenetic factor that plays a crucial role in gene regulation. It is a feature of different types of human diseases and is predominant in case of different cancer types. The epigenetic alterations have an effect on the genes participating in the tumorigenesis and metastasis of cancer (Nagarajan et al., 2009). In this study, UALCAN (Chandrashekar et al., 2017), which is an online web resource for the analysis of cancer OMICS data, was employed for obtaining the promoter methylation of hub genes through TCGA datasets for both GBM and HCC. The beta values indicated DNA methylation levels ranging from 0 (i.e. unmethylated) to 1 (i.e. fullymethylated). For hypermethylation, the specified range of beta value was 0.5 - 0.7 and for hypomethylation, this range was 0.05 - 0.3.

### Genetic Alterations of Hub Genes

The genetic alterations that mainly include mutations and DNA copy number alterations correspond to changes in the DNA sequences due to various factors. The accumulation of such genetic alterations may lead to cancer development, metastasis, growth, and resistance to therapy. This validation of genetic alterations in the hub genes was accomplished through cBioPortal (Gao et al., 2013), which is an open-source, open-access resource for interactively exploring the multidimensional cancer genomics data sets. For this purpose, 592 TCGA samples were considered for GBM and 391 samples for HCC. Copy number data sets were generated via GISTIC (Genomic Identification of Significant Targets in Cancer) algorithms that identify those regions that are significantly altered across the sets of patients. OncoPrints are used for visualization the genomic alterations (mutations and copy number alterations), and mRNA expression changes across a set of TCGA cases for the hub genes. In case of mutations, a splice site mutation occurs in an intronic region while splice region mutations takes place near the exon/intron junction. The copy number analysis derived from GISTIC algorithms indicates the level of copy-number per gene. In this case, -2 indicate deep deletion or deep loss and correspond to homozygous deletion. -1 corresponds to shallow deletion and indicates a heterozygous deletion. 0 is assigned to normal or diploid. 1 corresponds to gain that indicates low-level gain and 2 correspond to amplification and it indicates a high-level amplification.

### Differential Expression pattern validation and Survival Analysis of Hub Genes

The gene expression profiles of normal and cancerous TCGA samples related to all the 13 hub genes in both GBM and HCC were obtained through GEPIA (Gene Expression Profiling Interactive Analysis), an online web server (Tang et al., 2017). Thereafter, the survival analysis of these hub genes was obtained via web based tool, SurvExpress (Aguirre-Gamboa et al., 2013). The TCGA dataset in this case contained 148 patient samples of GBM and 361 patient samples of HCC. The univariate Cox regression analysis was employed to obtain the risk score by grouping the patients into high- and low-risk groups. Further, the Kaplan-Meier plot was obtained for visualizing the survival analysis of all the 13 hub genes (potential biomarkers) in both GBM and HCC.

## RESULTS

### Differentially Expressed Genes

There are a total of 3265 differentially expressed genes (1570 upregulated and 1695 downregulated) obtained from GBM datasets and 2321 differentially expressed genes (1444 upregulated and 877 downregulated) from HCC (see Supplementary Figure 1). The normal and cancerous tissues of brain (GBM) and liver (HCC) cancer are taken from Human Protein Atlas (HPA) (see Supplementary Figure 2). Out of these differentially expressed genes obtained for both GBM and HCC, there are 757 differentially expressed genes (452 upregulated and 305 downregulated) that are shared between both GBM and HCC. These 757 differentially expressed genes are considered for further analysis of network and pathway enrichment. These are common DEGs that are taken forward from the same NGS analyzed data.

### Protein-Protein Interaction Network Analysis

The PPI network for the differentially expressed genes contained 757 nodes and 6628 edges (see Supplementary Figure 3). The PPI enrichment p-value was < 1.0e-16. Since this value is small, it indicates that the nodes are not random and the observed number of edges is significant. The modules obtained from the MCODE plug-in and subsequently cytohubba plug-in provided 13 common hub genes in both GBM and HCC, viz Assembly Factor for Spindle Microtubules (ASPM), Aurora Kinase A (AURKA), BUB1 Mitotic Checkpoint Serine/Threonine Kinase (BUB1), BUB1 Mitotic Checkpoint Serine/Threonine Kinase B (BUB1B), Cyclin A2 (CCNA2), Cyclin B2 (CCNB2), Kinase Family Member 2C (KIF2C), Maternal Embryonic Leucine Zipper Kinase (MELK), Non-SMC Condensin I Complex Subunit G (NCAPG), Non-SMC Condensin I Complex Subunit H (NCAPH), NUF2 Component of NDC80 Kinetochore Complex (NUF2), PDZ Binding Kinase (PBK), and DNA Topoisomerase II Alpha (TOP2A) (see Figure 2). All these genes had a function associated with chromosome and spindle behavior of mitotic cell division and showed an up-regulated expression level in both GBM and HCC.

**Figure 2.**
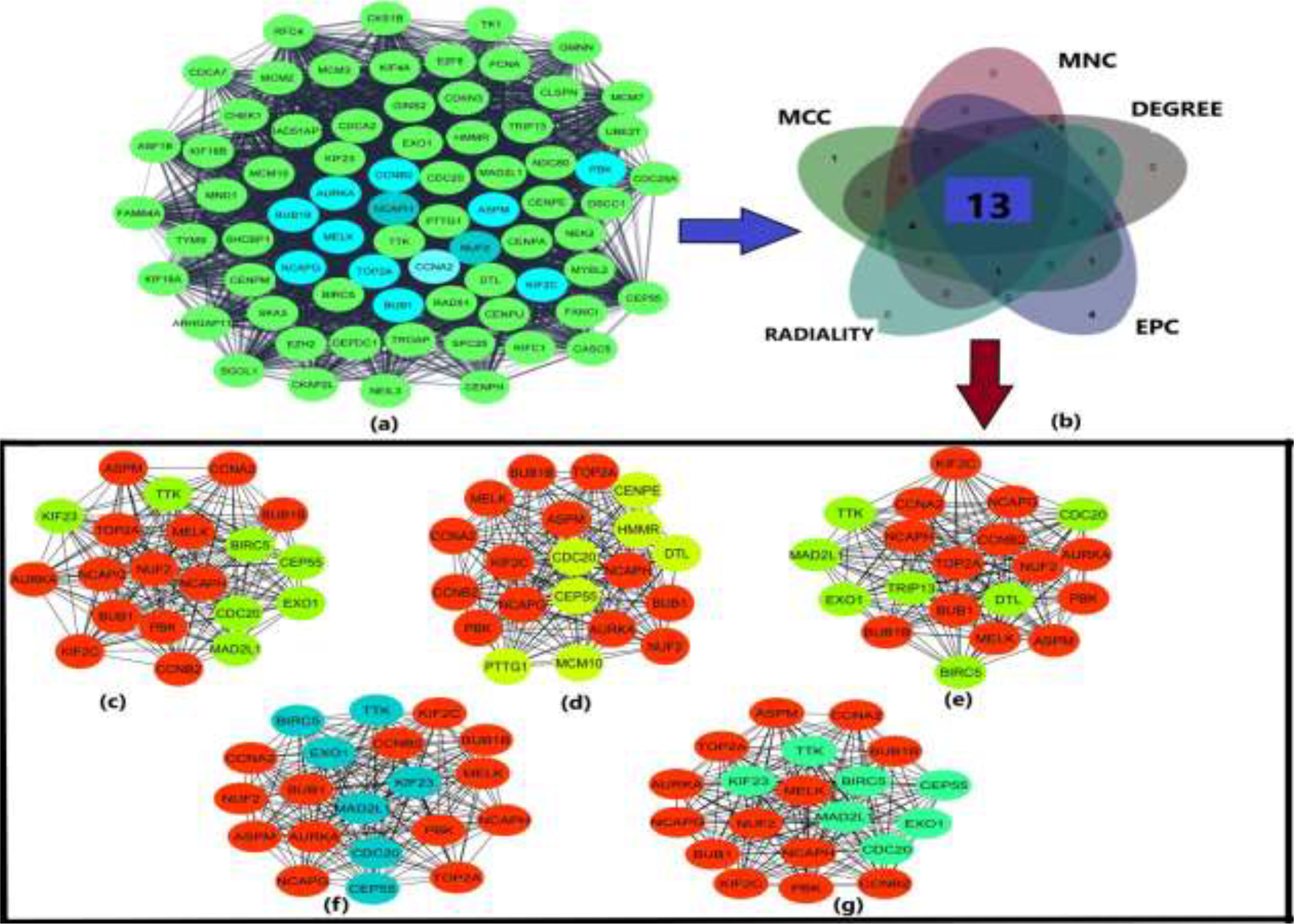
**(a)** Module of the PPI network obtained from MCODE plug-in of cytoscape **(b)** Venn diagram obtained from jvenn to find the common hub genes **(c)** 13 hub genes using degree method of cytohubba shown in red **(d) 13** hub genes using EPC method of cytohubba shown in red **(e)** 13 hub genes using MCC algorithm of cytohubba shown in red **(f)** 13 hub genes using MNC algorithm of cytohubba shown in red **(g)** 13 hub genes using radiality algorithm of cytohubba shown in red

The pairwise correlation analysis of DEGs using Pearson correlation statistics showed higher degree of positive correlation.

### GO Component and Pathway Enrichment Analysis

The results obtained for Biological processes from DAVID database showed that the hub genes are enriched in chromosome segregation, cell division, cell cycle process, nuclear division, and antigen processing and presentation. Likewise, the KEGG pathway analysis showed the involvement of 13 hub genes in cell cycle, DNA replication, oocyte meiosis, progesterone-mediated oocyte maturation, viral carcinogenesis, and Epstein-Barr virus infection signaling pathways (see Figure 3).

**Figure 3.**
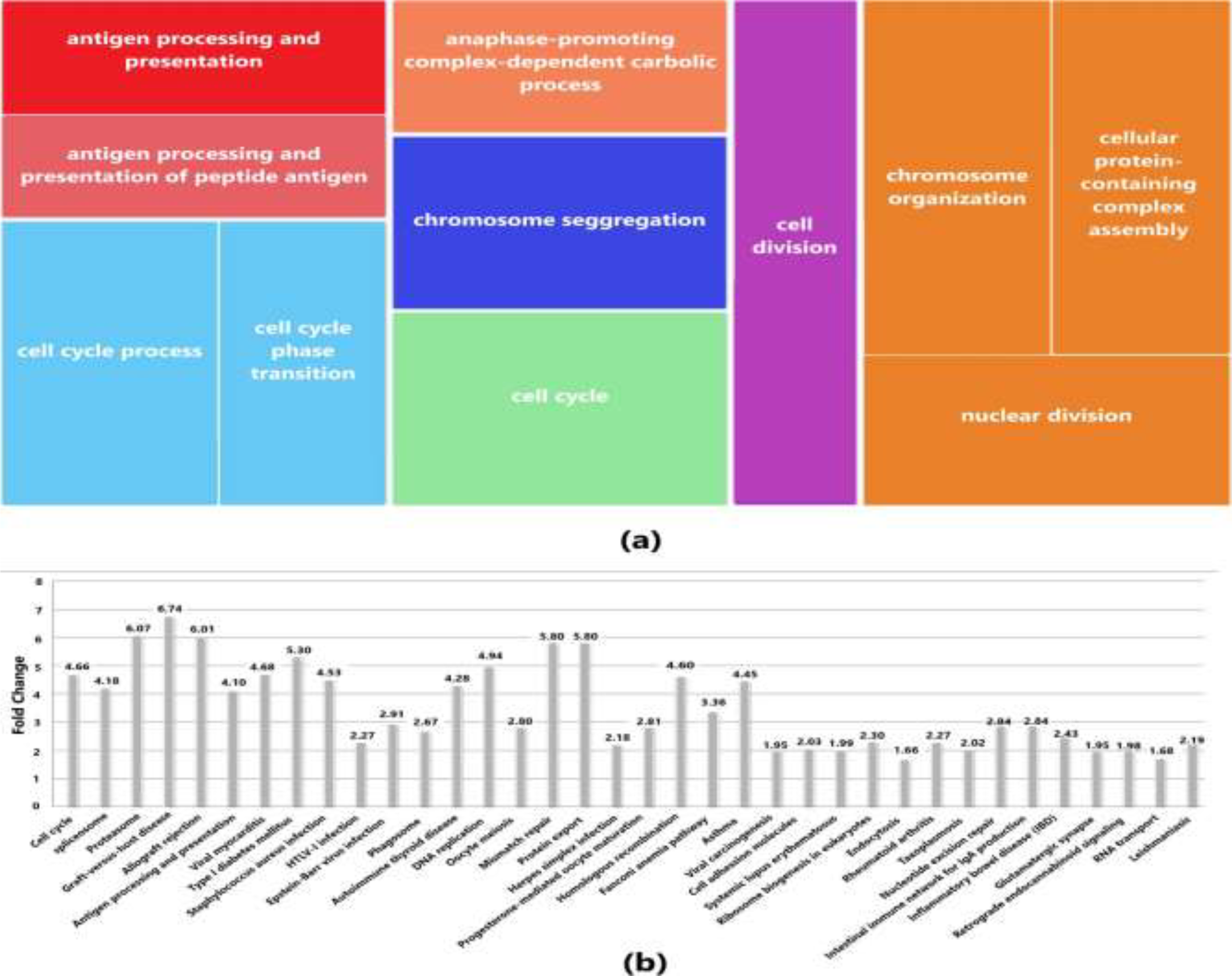
**(a)** Biological Processes (BP) based on p-values drawn from Revigo in which the hub genes are enriched **(b)** KEGG pathways corresponding to the enrichment of hub genes based on p-values and fold change

### Epigenetic Regulation of Gene Expression of Hub Genes by Promoter Methylation

Validation of promoter methylation using UALCAN database revealed that the promoter methylation level of ASPM, AURKA, BUB1, KIF2C, NCAPG, NCAPH, and NUF2 was lower than normal samples in GBM that indicates higher expression of these hub genes as against that of BUB1B, CCNA2, CCNB2, MELK, PBK, and TOP2A having higher promoter methylation level than normal samples (see Supplementary Figure 4(a)).

In case of HCC, the expression level of BUB1, CCNA2, CCNB2, KIF2C, MELK, NCAPG, NCAPH, NUF2, PBK, and TOP2A was higher due to their lower promoter methylation level against normal samples while ASPM, AURKA, and BUB1B were lowly expressed (see Supplementary Figure 4(b)).

### Differential Expression Pattern and Survival Analysis validation of Prognostic Biomarkers

The differential expression between normal and tumor cells obtained from GEPIA database showed that the expression of hub genes was significantly higher in case of GBM as compared to HCC. Moreover, among the 13 hub genes, the expression level of TOP2A gene was significantly higher in both GBM and HCC (see Supplementary Figure 5).

The aberrant expression of ASPM, AURKA, BUB1, BUB1B, MELK, NUF2, and PBK resulted in poorer survival rate of GBM patients in the high-risk group with survival rate less than 2 years. The median survival rate was less than 2 years for all the 13 hub genes (see Figure 4). For each patient, the risk score was calculated and ranking was done accordingly in the TCGA dataset. Patients were then divided into a high-risk group and a low-risk group.

**Figure 4.**
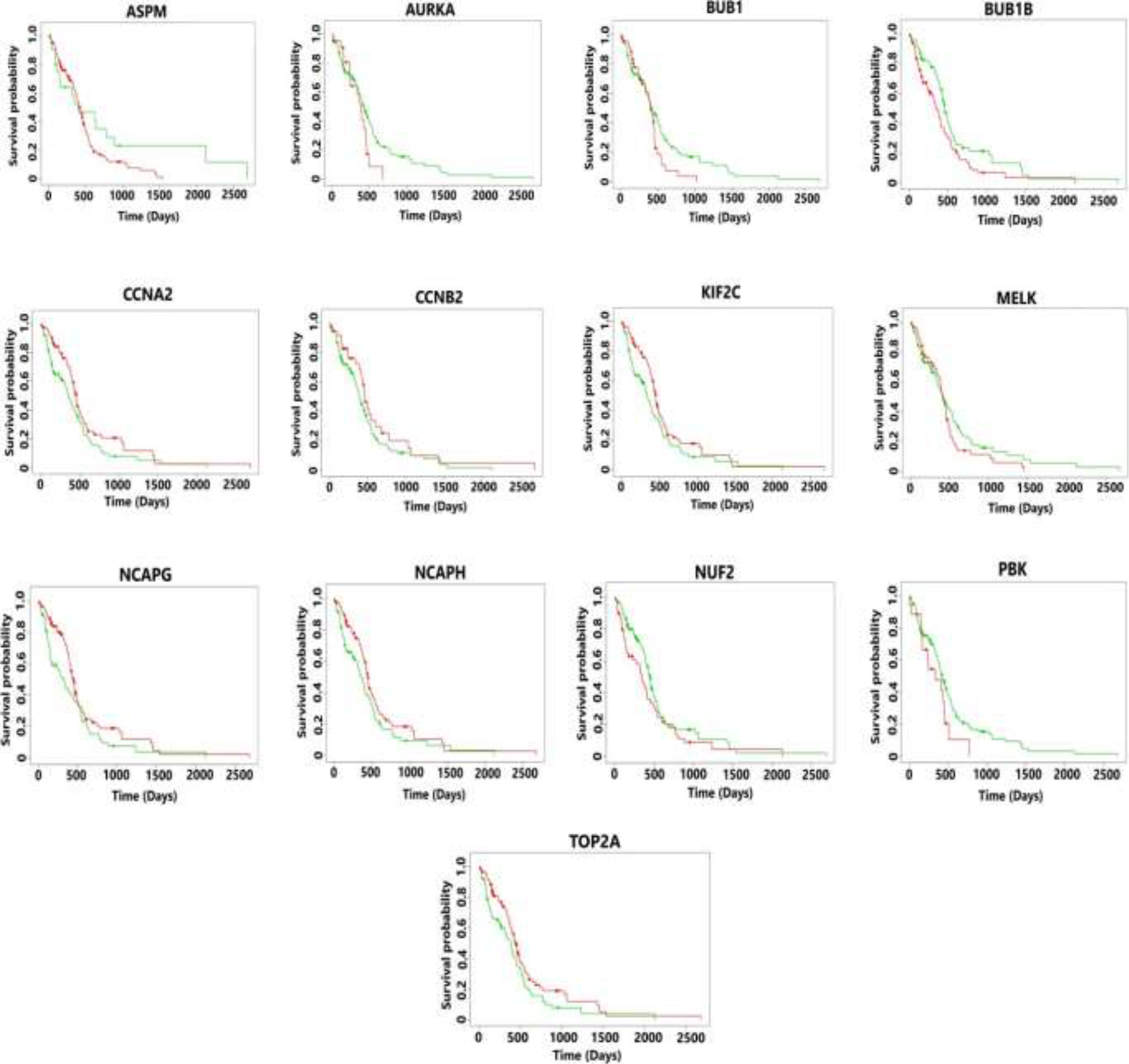
Kaplan-Meier plots showing the survival analysis of hub genes in GBM. The patients were divided into high- and low-risk groups. The overexpression of all the hub genes resulted in the poor survival outcomes which is less than 2 years for the patients suffering from GBM

Risk score (RS) = 0.03 * expression _ASPM_ + 0.041 * expression _AURKA_ + 0.044 * expression _BUB1_ – 0.03 * expression

_BUB1B_ + 0.011 * expression _CCNA2_ + 0.007 * expression _CCNB2_ + 0.046 * expression _KIF2C_ + 0.009 * expression _MELK_ +

0.01 * expression _NCAPG_ + 0.013 * expression _NCAPH_ – 0.032 * expression _NUF2_ + 0.108 * expression _PBK_ + 0.035 *expression _TOP2A_

The hazard ratio > 1 for these hub genes also showed the higher level of survival risk (see Table 1)

**Table 1.** Table showing the survival analysis results of hub genes in GBM

| *Gene* | *Cox coefficient* | *Hazard Ratio (CI)* |
| --- | --- | --- |
| ASPM | 0.030 | 1.58 (0.89 – 2.79) |
| AURKA | 0.041 | 1.57 (0.93 – 2.68) |
| BUB1 | 0.044 | 1.45 (0.96 – 2.21) |
| BUB1B | -0.030 | 1.52 (1.05 – 2.21) |
| CCNA2 | 0.011 | 0.69 (0.48 – 1.00) |
| CCNB2 | 0.007 | 0.73 (0.47 – 1.15) |
| KIF2C | 0.046 | 0.71 (0.49 – 1.02) |
| MELK | 0.009 | 1.23 (0.85 – 1.79) |
| NCAPG | 0.010 | 0.68 (0.47 – 0.99) |
| NCAPH | 0.013 | 0.73 (0.50 – 1.05) |
| NUF2 | -0.032 | 1.32 (0.91 – 1.93) |
| PBK | 0.108 | 1.64 (0.94 – 2.85) |
| TOP2A | 0.035 | 0.70 (0.49 – 1.01) |

The survival analysis of patients in the high-risk group showed poorer median survival rate which was less than 3 years (see Figure 5). The risk score was calculated as shown below-

**Figure 5.**
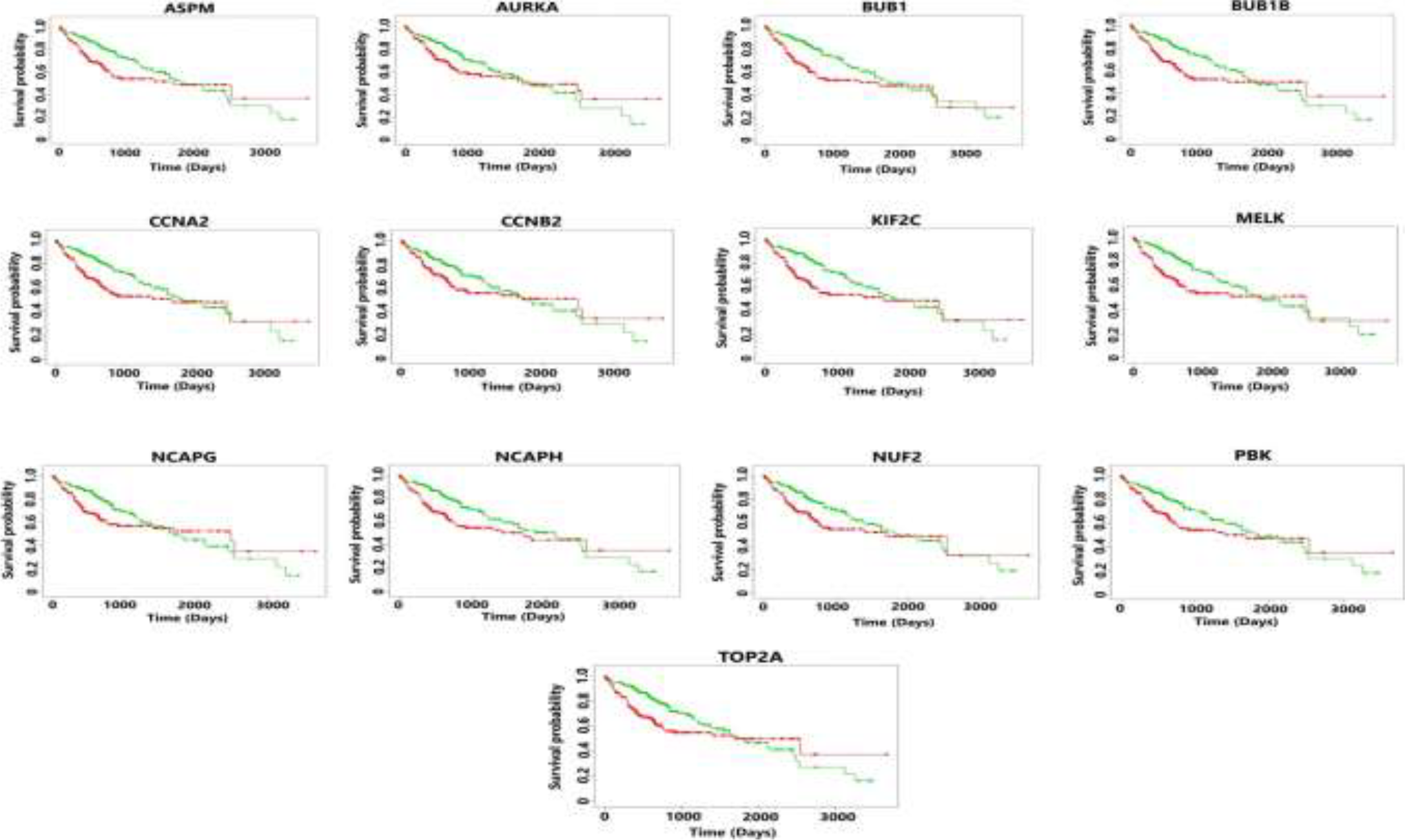
Kaplan-Meier plots showing the survival analysis corresponding to hub genes in HCC. The patients were divided into high- and low-risk groups. The overexpression of all the hub genes resulted in the poor survival outcomes which is less than 3 years for the patients suffering from metastatic HCC

Risk score (RS) = 0.078 * expression _ASPM_ + 0.02 * expression _AURKA_ + 0.25 * expression _BUB1_ + 0.268 * expression

_BUB1B_ + 0.173 * expression _CCNA2_ + 0.139 * expression _CCNB2_ + 0.278 * expression _KIF2C_ + 0.278 * expression _MELK_ +

0.153 * expression _NCAPG_ + 0.235 * expression _NCAPH_ + 0.268 * expression _NUF2_ + 0.233 * expression _PBK_ + 0.062 *expression _TOP2A_

The hazard ratio > 1 for all the 13 hub genes also indicated poorer survival rate of HCC patients (see Table 2).

**Table 2.** Table showing the survival analysis results of hub genes in HCC

| *Gene* | *Cox coefficient* | *Hazard Ratio (CI)* |
| --- | --- | --- |
| ASPM | 0.078 | 1.50 (1.05 – 2.13) |
| AURKA | 0.020 | 1.28 (0.90 – 1.81) |
| BUB1 | 0.250 | 1.68 (1.18 – 2.39) |
| BUB1B | 0.268 | 1.51 (1.06 – 2.14) |
| CCNA2 | 0.173 | 1.56 (1.10 – 2.21) |
| CCNB2 | 0.139 | 1.27 (0.90 – 1.80) |
| KIF2C | 0.278 | 1.57 (1.10 – 2.23) |
| MELK | 0.278 | 1.62 (1.14 – 2.30) |
| NCAPG | 0.153 | 1.31 (0.92 – 1.85) |
| NCAPH | 0.235 | 1.53 (1.08 – 2.18) |
| NUF2 | 0.268 | 1.48 (1.04 – 2.11) |
| PBK | 0.233 | 1.47 (1.03 – 2.08) |
| TOP2A | 0.062 | 1.44 (1.02 – 2.05) |

### Genetic Alterations in Hub Genes

Tumorigenesis mainly occurs due to irremediable mutations in cell structures. These mutations could be identified through genetic alteration analysis. The alterations may be in the form of missense mutation, splice mutation, deep deletion, truncating mutation, and amplification. In case of GBM, the percentage alteration of all the 13 hub genes varied from 0.3% to 2.1% (see Supplementary Figure 6). The corresponding copy number variations are shown in Supplementary Figure 7. The details of genetic alterations and copy number variations can be found in the table (see Supplementary Table 1). Most of the mutations in hub genes were found at phosphorylation, acetylation and uniquitination PTM sites with the characteristics of missense mutation and diploid copy type alteration.

In case of HCC, the alteration percentage had variations between 0.3% - 10% for the 13 prognostic biomarkers (see Supplementary Figure 8). The copy number variations are shown in Supplementary Figure 9. The description of genetic alterations and copy number variations are summarized in Supplementary Table 2. The results shows that mutations mainly occurred at phosphorylation and ubiquitination PTM sites with diploid copy number variations and missense mutations and these features were found to be enriched in the tumorigenesis and metastasis of cancers with markedly stronger accumulation and evolutionary conservation in protein domains (Narayan et al., 2016).

## Discussion

Cancer development due to uncontrolled cell division is the leading cause of death worldwide. The most dangerous event that leads to cancer development is mitosis having irreversible segregation of sister chromatids to daughter cells (Jallepalli et al., 2001). Abnormal chromosome segregation during mitosis results in tumorigenesis. This happens mainly due to failure in the mechanism of spindle assembly checkpoint as the checkpoint ensures proper chromosome segregation during mitosis (Baker et al., 2005). This chromosomal instability that results in abnormal chromosome numbers produces uncontrolled cell division leading to tumorigenesis and subsequently to metastasis of cancer types (Cahill et al., 1998). Hepatocellular carcinoma (HCC) is one of the leading cancer types that metastasizes to the lungs, adrenal glands, lymph nodes, and brain (Kummar et al., 2003). The evidence of brain metastasis from HCC is rare but is nowadays becoming more frequent compared to the conditions in the past (Wang et al., 2017). This study mainly focused on the metastasis of HCC to the brain that could potentially lead to the development of Glioblastoma Multiforme (GBM), the IV grade brain cancer. This progression and metastasis of this cancer resulted from the aberrant function of some genes and alteration in the patterns of gene expression. This dysregulation in the gene expression is mainly due to genetic alterations such as mutation, amplification, and copy number alterations (Jones et al., 2007). Thirteen hub genes were obtained from the protein-protein interaction network analysis using differentially expressed genes. Similarly, the pathway enrichment analysis done using DAVID database showed the involvement of these hub genes in processes like cell cycle, cell cycle process, and oocyte meiosis. These signaling pathways actively participate in cancer development, leading to tumorigenesis and metastasis.

Further validation of these genes was carried out using UALCAN database that found them to be hypomethylated in both HCC and GBM. This resulted in their aberrant expression through the increased probability of undergoing mutations leading to tumorigenesis (Wajed et al., 2001). All the 13 hub genes i.e. ASPM, AURKA, BUB1, BUB1B, CCNA2, CCNB2, KIF2C, MELK, NCAPG, NCAPH, NUF2, PBK, and TOP2A are oncogenes and were found to be upregulated in all the samples of HCC and GBM. ASPM gene had abnormality due to its overexpression in HCC and played a vital role in cell proliferation and metastasis (Lin et al., 2008). It promotes the progression of HCC through the activation of Wnt/β‐catenin signaling (Zhang et al., 2021). This gene had missense mutations at 8 different locations in 2% of the patients on phosphorylation and ubiquitination post-transcriptional modification (PTM) sites (refer to table 4). These mutations happened due to diploid copy number alteration. Likewise, in case of GBM, it had missense mutations on phosphorylation, acetylation, ubiquitination, and methylation PTM sites at 17 different locations (refer to table 3). Aurora Kinase A (AURKA) also has tumorigenesis properties in different cancer types (Du et al., 2021). This gene was involved in the cancer metastases in case of HCC (Chen et al., 2017). The missense mutation having diploid type alteration in 0.57% of the patients on phosphorylation PTM sites at 2 locations resulted in the abnormality in this gene. Likeweise, in case of GBM, this gene has missense mutations in 0.57% of the patients, with diploid copy number alterations on phosphorylation and acetylation PTM sites at 3 different locations. In one of the studies, it was found that AURKA inhibition suppressed the cell proliferation of GBM (Barton et al., 2010). BUB1 overexpression promoted tumorigenesis and aneuploidy (Ricke et al., 2011). This resulted in poorer survival of patients suffering from HCC (Yang et al. 2019). It had missense mutations in 0.57% of the patients at 2 different locations with diploid copy number alterations. In GBM also, upregulated BUB1 was also responsible for cell proliferation resulting in tumorigenesis (Yu et al., 2019). It has missense and splice mutations at 5 different locations. It has phosphorylation and ubiquitination PTM sites with diploid and shallow deletion type of copy number alterations. The next hub gene BUB1B was involved in progression of hepatocellular carcinoma (HCC) by activating mTORC1 signalling pathway (Qiu et al., 2020). It has missense mutation at a single location in 0.29% of the patients. It has diploid copy number alterations. In case of GBM, BUB1B was found to promote tumor proliferation (Ma et al., 2017). It has splice mutation in 0.29% of the patients with at a single location and has diploid copy number alterations associated with it. CCNA2 found to promote uncontrolled cell growth resulting in tumorigenesis in case of different cancer types (Gan et al., 2018; Lee et al., 2020; Gao et al., 2014). The upregulated CCNA2 was involved in cell cycle progression that resulted in tumorigenesis and metastasis in case of HCC (Gayard et al., 2018). It had missense mutation in 0.29% of the patients with diploid copy number alterations. In case of GBM, overexpressed CCNA2 resulted in poor prognosis of patients (Yang et al., 2020). It had missense and nonsense mutations in 0.29% of the patients on acetylation, phosphorylation and ubiquitination PTM sites on A25V and E269. CCNB2 also found to promote cell cycle progression resulting in tumorigenesis in case of triple negative breast cancer (Wu et al., 2021). It was also identified in the cell cycle progression leading to poor prognosis of HCC (Liu et al., 2020). In this study, amplification was found as genetic alterations in 0.57% of the patients. In GBM, CCNB2 acted as potential biomarker and played a vital role in Cellular Senescence and cell cycle (Jiang et al., 2020). These have missense mutation at location P80S having phosphorylation PTM site and amplification in 0.57% of the patients and shallow deletion copy number alterations. The absence of mutation in HCC and presence of one mutation in GBM showed that this mutation might have taken place due to metastasis of HCC in brain leading to GBM. KIF2C resulted in tumorigenesis due to abnormal cell cycle progression and metastasis in cervical cancer (Yang et al., 2022). In case of HCC, it participated in the progression of HCC and could be a potential therapeutic target (Gao et al., 2021). On the other hand, in case of GBM, it had missense mutations at three different locations on phosphorylation, ubiquitination, acetylation, and methylation PTM sites with diploid and gain copy number alterations. According to a study, this MELK gene was found to possess therapeutic drug like property due to its role in cell proliferation and triggering of cell cycle arrest in different cancer types (Giuliano et al., 2018). Its overexpression in case of HCC strongly correlated with abnormal cell growth leading to early recurrence and poor prognosis of patients (Xia et al., 2016). In the following study, it had missense mutations in 1.43% of the patients at 4 different locations on phosphorylation PTM sites having diploid copy number alterations. In GBM, MELK developed tumorigenesis and its inhibition could effectively suppress the abnormal growth of GBM (Zhang et al., 2020). Here, it was found to have missense mutations in 1.43% of the patients. It has phosphorylation PTM site and diploid copy number alterations. NCAPG gene was found to be responsible for survival of tumor cell leading to tumorigenesis and metastasis in HCC (Wang et al., 2019). It had missense and nonsense mutations in 0.86% of the patients at 3 different locations. It had diploid copy number alterations on phosphorylation and ubiquitination PTM sites. Similarly, NCAPG was responsible for promoting tumor progression in case of GBM also (Zheng et al., 2022). It had missense, splice and nonsense type mutations in 0.86% of the patients at 5 different locations on phosphorylation, ubiquitination PTM sites and diploid, gain and shallow deletion copy number alterations. NCAPH was found to be overexpressed in different cancer types promoting tumorigenesis and possibly metastasis (Qiu et al., 2020; Cui et al., 2019; Yin et al., 2017). The upregulation of NCAPH resulted in the enhancement of cell proliferation, invasion and migration in case of HCC (Sun et al., 2019). In this study, it had missense mutations in 0.29% of the patients having diploid copy number alterations. In case of GBM, it had missense mutations at in 0.29% of the patients with diploid copy number alterations. This gene played a regulatory role in the cell proliferation and apoptosis in case of HCC (Liu et al., 2019). It had missense mutation at 3 locations in 0.86% of the patients having gain type of copy number alterations. In GBM, NUF2 promoted tumorigenesis and its downregulation had inhibited growth of tumor cells and induced apoptosis (Hu et al., 2015). It had missense mutation in 0.79% of the patients at three different locations on phosphorylation and ubiquitination PTM sites having diploid copy number alterations. The overexpression of PBK in case of HCC promoted metastasis through the activation of ETV4-uPAR signaling pathway (Yang et al., 2019). It had missense mutation at E303V in 0.29% of the patients having diploid copy number alterations in HCC. Similarly, its overexpression resulted in poorer survival rate in case of GBM (Dong et al., 2020). It had missense mutations in about 0.26% of the patients on phosphorylation PTM sites having diploid copy number alterations. TOP2A was associated with growth in HCC tumor cells resulting in metastasis (Cai et al., 2020). It had nonsense and missense mutations in 1.14% of the patients at 4 different locations on phosphorylation PTM sites having diploid and gain copy number alterations. In case of GBM also, TOP2A had missense mutations at 5 different locations in 0.79% of the patients on phosphorylation, sumoylation, acetylation, ubiquitination, and methylation PTM sites having diploid copy number alterations. These 13 hub genes that were discussed above were associated with worst survival of the patients as studied through Kaplan-Meier survival plots in case of both GBM and HCC. This survival rate was less than 2 years due to overexpression of these genes and hence these could be potential prognostic biomarkers that could help in the suppression of metastasis of HCC.

## Conclusion

The present study identified 13-gene signature i.e. ASPM, AURKA, BUB1, BUB1B, CCNA2, CCNB2, KIF2C, MELK, NCAPG, NCAPH, NUF2, PBK, and TOP2A. These 13 hub genes could behave as potential biomarkers as their overexpression resulted in the abnormal cell division leading to tumorigenesis and metastasis in HCC and this cancer metastasized in the brain causing GBM. This overexpression resulted in the poor survival of patients in both GBM and HCC. Proper design of suitable inhibitors for these overexpressed hub genes will help in reducing the tumorigenesis and metastasis of HCC thereby increasing the overall survival outcomes of the patients.

## Supporting information

supplementary images

## References

Ardito, F., Giuliani, M., Perrone, D., Troiano, G., & Muzio, L. Lo. (2017) ‘The crucial role of protein phosphorylation in cell signaling and its use as targeted therapy (Review)’. International Journal of Molecular Medicine, vol. 40 No. 2, pp.271–280. https://doi.org/10.3892/ijmm.2017.3036

Baker, D. J., Chen, J., & Van Deursen, J. M. A. (2005) ‘The mitotic checkpoint in cancer and aging: What have mice taught us?’ Current Opinion in Cell Biology, vol. 17 No. 6, pp. 583–589. https://doi.org/10.1016/j.ceb.2005.09.011

Bardou, P., Mariette, J., Escudié, F., Djemiel, C., & Klopp, C. (2014) ‘SOFTWARE Open Access jvenn: an interactive Venn diagram viewer’. BMC Bioinformatics, vol. 15 No. 293, pp. 1–7. http://www.biomedcentral.com/1471-2105/15/293

Barton, V. N., Foreman, N. K., Donson, A. M., Birks, D. K., Handler, M. H., & Vibhakar, R. (2010) ‘Aurora kinase A as a rational target for therapy in glioblastoma: Laboratory investigation’. Journal of Neurosurgery: Pediatrics, vol. 6 No. 1, pp. 98–105. https://doi.org/10.3171/2010.3.PEDS10120

Bayard, Q., Meunier, L., Peneau, C., Renault, V., Shinde, J., Nault, J. C., Mami, I., Couchy, G., Amaddeo, G., Tubacher, E., Bacq, D., Meyer, V., La Bella, T., Debaillon-Vesque, A., Bioulac-Sage, P., Seror, O., Blanc, J. F., Calderaro, J., Deleuze, J. F., &Letouzé, E. (2018) ‘Cyclin A2/E1 activation defines a hepatocellular carcinoma subclass with a rearrangement signature of replication stress’. Nature Communications, vol. 9 No. 1. https://doi.org/10.1038/s41467-018-07552-9

Brinton, L. T., Brentnall, T. A., Smith, J. A., & Kelly, K. A. (2012) ‘Metastatic biomarker discovery through proteomics’. Cancer Genomics and Proteomics, vol. 9 No. 6, pp. 345–356.

Cahill, D. P., Lengauer, C., Yu, J., Riggins, G. J., Willson, J. K. V., Markowitz, S. D., Kinzler, K. W., & Vogelstein, B. (1998) ‘Mutations of mitotic checkpoint genes in human cancers’. Nature, vol. 392 No. 6673, pp. 300–303. https://doi.org/10.1038/32688

Cai, H., Shao, B., Zhou, Y., & Chen, Z. (2020) ‘High expression of TOP2A in hepatocellular carcinoma is associated with disease progression and poor prognosis’. Oncology Letters, vol. 20 No. 5, pp. 1–9. https://doi.org/10.3892/ol.2020.12095

Cao, Y. (2017) ‘Tumorigenesis as a process of gradual loss of original cell identity and gain of properties of neural precursor/progenitor cells’. Cell and Bioscience, vol. 7 No. 1, pp. 1–14. https://doi.org/10.1186/s13578-017-0188-9

Chakravarthi, B. V. S. K., Nepal, S., & Varambally, S. (2016) ‘Genomic and Epigenomic Alterations in Cancer’. American Journal of Pathology, vol. 186 No. 7, pp. 1724–1735. https://doi.org/10.1016/j.ajpath.2016.02.023

Chandrashekar, D. S., Bashel, B., Balasubramanya, S. A. H., Creighton, C. J., Ponce-Rodriguez, I., Chakravarthi, B. V. S. K., & Varambally, S. (2017) ‘UALCAN: A Portal for Facilitating Tumor Subgroup Gene Expression and Survival Analyses’. Neoplasia (United States), vol. 19 No. 8, pp. 649–658. https://doi.org/10.1016/j.neo.2017.05.002

Chen, C., Song, G., Xiang, J., Zhang, H., Zhao, S., & Zhan, Y. (2017) ‘AURKA promotes cancer metastasis by regulating epithelial-mesenchymal transition and cancer stem cell properties in hepatocellular carcinoma’. Biochemical and Biophysical Research Communications, vol. 486 No. 2, pp.514–520. https://doi.org/10.1016/j.bbrc.2017.03.075

Chen, M., Li, S., Liang, Y., Zhang, Y., Luo, D., & Wang, W. (2021) ‘Integrative Multi-Omics Analysis of Identified NUF2 as a Candidate Oncogene Correlates With Poor Prognosis and Immune Infiltration in Non-Small Cell Lung Cancer’. Frontiers in Oncology, vol. 11(June), pp. 1–13. https://doi.org/10.3389/fonc.2021.656509

Court, C. M., Hou, S., Liu, L., Winograd, P., DiPardo, B. J., Liu, S. X., Chen, P. J., Zhu, Y., Smalley, M., Zhang, R., Sadeghi, S., Finn, R. S., Kaldas, F. M., Busuttil, R. W., Zhou, X. J., Tseng, H. R., Tomlinson, J. S., Graeber, T. G., & Agopian, V. G. (2020) ‘Somatic copy number profiling from hepatocellular carcinoma circulating tumor cells’. Npj Precision Oncology, vol. 4 No. 1. https://doi.org/10.1038/s41698-020-0123-0

Cui, F., Hu, J., Xu, Z., Tan, J., & Tang, H. (2019) ‘Overexpression of ncaph is upregulated and predicts a poor prognosis in prostate cancer’. Oncology Letters, vol. 17 No. 6, pp. 5768–5776. https://doi.org/10.3892/ol.2019.10260

Dawood, S. (2010) ‘Novel biomarkers of metastatic cancer’. Expert Review of Molecular Diagnostics, vol. 10 No. 5, pp. 581–590. https://doi.org/10.1586/erm.10.35

Deng, L., Meng, T., Chen, L., Wei, W., & Wang, P. (2020) ‘The role of ubiquitination in tumorigenesis and targeted drug discovery’. Signal Transduction and Targeted Therapy, vol. 5 No. 1. https://doi.org/10.1038/s41392-020-0107-0

Dennis, G., Sherman, B. T., Hosack, D. A., Yang, J., Gao, W., Lane, H. C., & Lempicki, R. A. (2003) ‘DAVID: Database for Annotation, Visualization, and Integrated Discovery’. Genome Biology, vol. 4 No. 5. https://doi.org/10.1186/gb-2003-4-9-r60

Dong, C., Fan, W., & Fang, S. (2020) ‘PBK as a Potential Biomarker Associated with Prognosis of Glioblastoma’. Journal of Molecular Neuroscience, vol. 70 No. 1, pp. 56–64. https://doi.org/10.1007/s12031-019-01400-1

Du, R., Huang, C., Liu, K., Li, X., & Dong, Z. (2021) ‘Targeting AURKA in Cancer: molecular mechanisms and opportunities for Cancer therapy’. Molecular Cancer, vol. 20 No 1, pp. 1–27. https://doi.org/10.1186/s12943-020-01305-3

Ehrlich, M. (2002) ‘DNA methylation in cancer: Too much, but also too little’. Oncogene, vol. 21(35 REV. ISS. 3), pp. 5400–5413. https://doi.org/10.1038/sj.onc.1205651

Ehrlich, M., Woods, C. B., Yu, M. C., Dubeau, L., Yang, F., Weisenberger, D. J., Long, T. I., Youn, B., Emerich, S., & Laird, P. W. (2006) ‘NIH Public Access’. Vol. 25 No. 18, pp. 2636–2645.

Fares, J., Fares, M. Y., Khachfe, H. H., Salhab, H. A., & Fares, Y. (2020) ‘Molecular principles of metastasis: a hallmark of cancer revisited’. Signal Transduction and Targeted Therapy, vol. 5 No. 1. https://doi.org/10.1038/s41392-020-0134-x

Fasching, P. A., Brucker, S. Y., Fehm, T. N., Overkamp, F., Janni, W., Wallwiener, M., Hadji, P., Belleville, E., Häberle, L., Taran, F. A., Lüftner, D., Lux, M. P., Ettl, J., Müller, V., Tesch, H., Wallwiener, D., & Schneeweiss, A. (2015) ‘Biomarkers in patients with metastatic breast cancer and the praegnant study network’. Geburtshilfe Und Frauenheilkunde, vol. 75 No. 1, pp. 41–50. https://doi.org/10.1055/s-0034-1396215

Gan, Y., Li, Y., Li, T., Shu, G., & Yin, G. (2018) ‘CCNA2 acts as a novel biomarker in regulating the growth and apoptosis of colorectal cancer’. Cancer Management and Research, vol. 10, pp. 5113–5124. https://doi.org/10.2147/CMAR.S176833

Gao, J., Aksoy, B. A., Dogrusoz, U., Dresdner, G., Gross, B., Sumer, S. O., Sun, Y., Jacobsen, A., Sinha, R., Larsson, E., Cerami, E., Sander, C., & Schultz, N. (2014) ‘Integrative Analysis of Complex Cancer Genomics and Clinical Profiles Using the cBioPortal Complementary Data Sources and Analysis Options’. Science Signaling, vol. 6 No. 269, pp. 1–20. https://doi.org/10.1126/scisignal.2004088.

Gao, S., Zhu, D., Zhu, J., Shen, L., Zhu, M., & Ren, X. (2021) ‘Screening Hub Genes of Hepatocellular Carcinoma Based on Public Databases’. Computational and Mathematical Methods in Medicine, https://doi.org/10.1155/2021/7029130

Gao, T., Han, Y., Yu, L., Ao, S., Li, Z., & Ji, A. (2014) ‘CCNA2 is a prognostic biomarker for ER+ breast cancer and tamoxifen resistance’. PLoS ONE, vol. 9 No. 3, pp.1–5. https://doi.org/10.1371/journal.pone.0091771

Gao, Z., Jia, H., Yu, F., Guo, H., & Li, B. (2021) ‘KIF2C promotes the proliferation of hepatocellular carcinoma cells in vitro and in vivo’. Experimental and Therapeutic Medicine, vol. 22 No. 4, pp. 1–9. https://doi.org/10.3892/etm.2021.10528

Giuliano, C. J., Lin, A., Smith, J. C., Palladino, A. C., & Sheltzer, J. M. (2018) ‘MELK expression correlates with tumor mitotic activity but is not required for cancer growth’. ELife, vol. 7, pp. 1–23. https://doi.org/10.7554/eLife.32838

Hoffmann, M. J., & Schulz, W. A. (2005) ‘Causes and consequences of DNA hypomethylation in human cancer’. Biochemistry and Cell Biology, vol. 83 No. 3, pp. 296–321. https://doi.org/10.1139/o05-036

Hu, P., Shangguan, J., & Zhang, L. (2015) ‘Downregulation of NUF2 inhibits tumor growth and induces apoptosis by regulating lncRNA AF339813’. International Journal of Clinical and Experimental Pathology, vol. 8 No. 3, pp. 2638–2648.

Jallepalli, P. V., & Lengauer, C. (2001). Chromosome segregation and cancer: Cutting through the mystery. Nature Reviews Cancer, vol. 1 No. 2, pp. 109–117. https://doi.org/10.1038/35101065

Jiang, L., Zhong, M., Chen, T., Zhu, X., Yang, H., & Lv, K. (2020). Gene regulation network analysis reveals core genes associated with survival in glioblastoma multiforme. Journal of Cellular and Molecular Medicine, vol. 24 No. 17, pp. 10075–10087. https://doi.org/10.1111/jcmm.15615

Kamburov, A., Lawrence, M. S., Polak, P., Leshchiner, I., Lage, K., Golub, T. R., Lander, E. S., & Getz, G. (2015) ‘Comprehensive assessment of cancer missense mutation clustering in protein structures’. Proceedings of the National Academy of Sciences of the United States of America, vol. 112 No. 40, pp. E5486–E5495. https://doi.org/10.1073/pnas.1516373112

Kim, T. M., Yim, S. H., Shin, S. H., Xu, H. D., Jung, Y. C., Park, C. K., Choi, J. Y., Park, W. S., Kwon, M. S., Fiegler, H., Carter, N. P., Rhyu, M. G., & Chung, Y. J. (2008) ‘Clinical implication of recurrent copy number alterations in hepatocellular carcinoma and putative oncogenes in recurrent gains on 1q’. International Journal of Cancer, vol. 123 No. 12, pp.2808–2815. https://doi.org/10.1002/ijc.23901

Kummar, S., & Shafi, N. Q. (2003) ‘Metastatic hepatocellular carcinoma’. Clinical Oncology, vol. 15 No. 5, pp. 288–294. https://doi.org/10.1016/S0936-6555(03)00067-0

Lah, T. T., Novak, M., & Breznik, B. (2020) ‘Brain malignancies: Glioblastoma and brain metastases’. Seminars in Cancer Biology, vol. 60, pp. 262–273. https://doi.org/10.1016/j.semcancer.2019.10.010

Lee, Y., Lee, C. E., Oh, S., Kim, H., Lee, J., Kim, S. B., & Kim, H. S. (2020). Pharmacogenomic analysis reveals CCNA2 as a predictive biomarker of sensitivity to polo-like kinase i inhibitor in gastric cancer. Cancers, vol. 12 No. 6, pp. 1–14. https://doi.org/10.3390/cancers12061418

Li, Z., Lin, Y., Cheng, B., Zhang, Q., & Cai, Y. (2021) ‘Identification and Analysis of Potential Key Genes Associated With Hepatocellular Carcinoma Based on Integrated Bioinformatics Methods’. Frontiers in Genetics, vol. 12(March), pp. 1–14. https://doi.org/10.3389/fgene.2021.571231

Lin, S. Y., Pan, H. W., Liu, S. H., Jeng, Y. M., Hu, F. C., Peng, S. Y., Lai, P. L., & Hsu, H. C. (2008). ASPM\s a novel marker for vascular invasion, early recurrence, and poor prognosis of hepatocellular carcinoma. Clinical Cancer Research, vol. 14 No. 15, pp. 4814. https://doi.org/10.1158/1078-0432.CCR-07-5262

Liu, J., Peng, Y., & Wei, W. (2022) ‘Cell cycle on the crossroad of tumorigenesis and cancer therapy’. Trends in Cell Biology, vol. 32 No. 1, pp. 30–44. https://doi.org/10.1016/j.tcb.2021.07.001

Liu, J., Sun, G. L., Pan, S. L., Qin M. Bin, Ouyang, R., & Huang, J. A. (2020) ‘Identification of hub genes in colon cancer via bioinformatics analysis’. Journal of International Medical Research, vol. 48 No. 9. https://doi.org/10.1177/0300060520953234

Liu, L., Chen, A., Chen, S., Song, W., Yao, Q., Wang, P., & Zhou, S. (2020) ‘CCNB2, NUSAP1 and TK1 are associated with the prognosis and progression of hepatocellular carcinoma, as revealed by co-expression analysis’. Experimental and Therapeutic Medicine, pp. 2679–2689. https://doi.org/10.3892/etm.2020.8522

Lu, Y., Hu, J. G., Lin, X. J., & Li, X. G. (2017) ‘Bone metastases from hepatocellular carcinoma: clinical features and prognostic factors’. Hepatobiliary and Pancreatic Diseases International, vol. 16 No. 5, pp. 499–505. https://doi.org/10.1016/S1499-3872(16)60173-X

Ma, Q., Liu, Y., Shang, L., Yu, J., & Qu, Q. (2017) ‘The FOXM1/BUB1B signaling pathway is essential for the tumorigenicity and radioresistance of glioblastoma’. Oncology Reports, vol. 38 No. 6, pp. 3367–3375. https://doi.org/10.3892/or.2017.6032

Mo, S., Fang, D., Zhao, S., Thai Hoa, P. T., Zhou, C., Liang, T., He, Y., Yu, T., Chen, Y., Qin, W., Han, Q., Su, H., Zhu, G., Luo, X., Peng, T., & Han, C. (2022) ‘Down regulated oncogene KIF2C inhibits growth, invasion, and metastasis of hepatocellular carcinoma through the Ras/MAPK signaling pathway and epithelial-to-mesenchymal transition’. Annals of Translational Medicine, vol. 10 No. 3, pp.151–151. https://doi.org/10.21037/atm-21-6240

Narayan, S., Bader, G. D., & Reimand, J. (2016) ‘Frequent mutations in acetylation and ubiquitination sites suggest novel driver mechanisms of cancer’. Genome Medicine, vol. 8 No. 1, pp. 1–13. https://doi.org/10.1186/s13073-016-0311-2

Oura, K., Fujita, K., Morishita, A., Iwama, H., Nakahara, M. A. I., Tadokoro, T., Sakamoto, T., Nomura, T., Yoneyama, H., Mimura, S., Tani, J., Kobara, H., Okano, K., Suzuki, Y., & Masaki, T. (2019) ‘Serum microRNA-125a-5p as a potential biomarker of HCV-associated hepatocellular carcinoma’. Oncology Letters, vol. 18 No. 1, pp. 882–890. https://doi.org/10.3892/ol.2019.10385

Park, Y., Heider, D., & Hauschild, A. C. (2021) ‘Integrative analysis of next-generation sequencing for next-generation cancer research toward artificial intelligence’. Cancers, vol. 13 No. 13, pp. 1–20. https://doi.org/10.3390/cancers13133148

Paul Shannon, 1, Andrew Markiel, 1, Owen Ozier, 2 Nitin S. Baliga, 1 Jonathan T. Wang, 2 Daniel Ramage, 2, Nada Amin, 2, Benno Schwikowski, 1, 5 and Trey Ideker2, 3, 4, 5, 山本 隆久, 豊田 直平, 深瀬 吉邦, & 大森 敏行. (1971) ‘Cytoscape: A Software Environment for Integrated Models’. Genome Research, vol. 13 No. 22, pp. 426. https://doi.org/10.1101/gr.1239303.metabolite

Peter, A. J., & Baylin, S. B. (2007) ‘The Epigenomic of Cancer’. Cell, vol. 128 No 4, 683–692. https://doi.org/10.1016/j.cell.2007.01.029.The

Qiu, J., Zhang, S., Wang, P., Wang, H., Sha, B., Peng, H., Ju, Z., Rao, J., & Lu, L. (2020) ‘BUB1B promotes hepatocellular carcinoma progression via activation of the mTORC1 signaling pathway’. Cancer Medicine, vol. 9 No. 21, pp.8159–8172. https://doi.org/10.1002/cam4.3411

Qiu, X., Gao, Z., Shao, J., & Li, H. (2020) ‘NCAPH is upregulated in endometrial cancer and associated with poor clinicopathologic characteristics’. Annals of Human Genetics, vol. 84 No. 6, pp. 437–446. https://doi.org/10.1111/ahg.12398

Qu, G. Z., Dubeau, L., Narayan, A., Yu, M. C., & Ehrlich, M. (1999) ‘Satellite DNA hypomethylation vs. overall genomic hypomethylation in ovarian epithelial tumors of different malignant potential’. Mutation Research - Fundamental and Molecular Mechanisms of Mutagenesis, vol. 423 No. (1–2), pp. 91–101. https://doi.org/10.1016/S0027-5107(98)00229-2

Review, I. (2008) ‘Immune activation and inflammation in HIV-1 infection : June’, pp.231–241. https://doi.org/10.1002/path

Ricke, R. M., Jeganathan, K. B., & van Deursen, J. M. (2011) ‘Bub1 overexpression induces aneuploidy and tumor formation through Aurora B kinase hyperactivation’. Journal of Cell Biology, vol. 193 No. 6, pp.1049–1064. https://doi.org/10.1083/jcb.201012035

Riehn, M., Klopocki, E., Molkentin, M., Reinhardt, R., & Burmeister, T. (2011) ‘A BACH2-BCL2L1 Fusion Gene Resulting from a Lymphoma Cell Line BLUE-1’. Cancer, vol. 396(January), pp. 389–396. https://doi.org/10.1002/gcc

Roman-Gomez, J., Jimenez-Velasco, A., Agirre, X., Castillejo, J. A., Navarro, G., San Jose-Eneriz, E., Garate, L., Cordeu, L., Cervantes, F., Prosper, F., Heiniger, A., & Torres, A. (2008) ‘Repetitive DNA hypomethylation in the advanced phase of chronic myeloid leukemia’. Leukemia Research, vol. 32 No. 3, pp.487–490. https://doi.org/10.1016/j.leukres.2007.07.021

Scott, A., & Salgia, R. (2009) ‘Therapeutics and Decision Making’. Vol. 2 No. 6, pp. 577–586. https://doi.org/10.2217/17520363.2.6.577.Biomarkers

Sun, C., Huang, S., Wang, H., Xie, R., Zhang, L., Zhou, Q., He, X., & Ju, W. (2019) ‘Non-SMC condensin I complex subunit H enhances proliferation, migration, and invasion of hepatocellular carcinoma’. Molecular Carcinogenesis, vol. 58 No. 12, pp.2266–2275. https://doi.org/10.1002/mc.23114

Supek, F., Bošnjak, M., Škunca, N., & Šmuc, T. (2011) ‘Revigo summarizes and visualizes long lists of gene ontology terms’. PLoS ONE, vol. 6 No. 7. https://doi.org/10.1371/journal.pone.0021800

Szklarczyk, D., Gable, A. L., Nastou, K. C., Lyon, D., Kirsch, R., Pyysalo, S., Doncheva, N. T., Legeay, M., Fang, T., Bork, P., Jensen, L. J., & von Mering, C. (2021) ‘The STRING database in 2021: Customizable protein-protein networks, and functional characterization of user-uploaded gene/measurement sets’. Nucleic Acids Research, vol. 49 No. D1, pp. D605–D612. https://doi.org/10.1093/nar/gkaa1074

Tang, Y., Zhang, Y., & Hu, X. (2020) ‘Identification of Potential Hub Genes Related to Diagnosis and Prognosis of Hepatitis B Virus-Related Hepatocellular Carcinoma via Integrated Bioinformatics Analysis’. BioMed Research International, https://doi.org/10.1155/2020/4251761

Tang, Z., Li, C., Kang, B., Gao, G., Li, C., & Zhang, Z. (2017) ‘GEPIA: A web server for cancer and normal gene expression profiling and interactive analyses’. Nucleic Acids Research, vol. 45 No. W1, pp. W98–W102. https://doi.org/10.1093/nar/gkx247

Testa, U., Castelli, G., & Pelosi, E. (2020) ‘Genetic alterations of metastatic colorectal cancer’. Biomedicines, vol. 8 No. 10, pp. 1–29. https://doi.org/10.3390/biomedicines8100414

Theivendran, S., Tang, J., Lei, C., Yang, Y., Song, H., Gu, Z., Wang, Y., Yang, Y., Jin, L., & Yu, C. (2020) ‘Post translational modification-assisted cancer immunotherapy for effective breast cancer treatment’. Chemical Science, vol. 11 No. 38, pp. 10421–10430. https://doi.org/10.1039/d0sc02803g

Tischoff, I., & Tannapfel, A. (2008) ‘DNA methylation in hepatocellular carcinoma’. World Journal of Gastroenterology, vol. 14 No. 11, pp.1741–1748. https://doi.org/10.3748/wjg.14.1741

Wajed, S. A., Laird, P. W., & DeMeester, T. R. (2001) ‘DNA methylation: An alternative pathway to cancer’. Annals of Surgery, vol. 234 No. 1, pp.10–20. https://doi.org/10.1097/00000658-200107000-00003

Wang, S., Wang, A., Lin, J., Xie, Y., Wu, L., Huang, H., Bian, J., Yang, X., Wan, X., Zhao, H., & Huang, J. (2017) ‘Brain metastases from hepatocellular carcinoma: Recent advances and future avenues’. Oncotarget, vol. 8 No. 15, pp.25814–25829. https://doi.org/10.18632/oncotarget.15730

Wang, Y., Gao, B., Tan, P. Y., Handoko, Y. A., Sekar, K., Deivasigamani, A., Seshachalam, V. P., Ouyang, H. Y., Shi, M., Xie, C., Goh, B. K. P., Ooi, L. L., & Hui, K. M. (2019) ‘Genome-wide CRISPR knockout screens identify NCAPG as an essential oncogene for hepatocellular carcinoma tumor growth’. FASEB Journal, vol. 33 No. 8, pp. 8759–8770. https://doi.org/10.1096/fj.201802213RR

Wang, Y., Sibaii, F., Lee, K. J. Gill, M., & L. Hatch, J. (2021) ‘NOTE: This preprint reports new research that has not been certified by peer review and should not be used to guide clinical practice’. Vol. 1. MedRxiv No. 165, pp. 1–13.

Wu, S., Su, R., & Jia, H. (2021) ‘Cyclin B2 (CCNB2) Stimulates the Proliferation of Triple-Negative Breast Cancer (TNBC) Cells in Vitro and in Vivo’. Disease Markers. https://doi.org/10.1155/2021/5511041

Xia, H., Kong, S. N., Chen, J., Shi, M., Sekar, K., Seshachalam, V. P., Rajasekaran, M., Goh, B. K. P., Ooi, L. L., & Hui, K. M. (2016) ‘MELK is an oncogenic kinase essential for early hepatocellular carcinoma recurrence’. Cancer Letters, vol. 383 No. 1, pp. 85–93. https://doi.org/10.1016/j.canlet.2016.09.017

Yang, J., Wu, Z., Yang, L., Jeong, J. H., Zhu, Y., Lu, J., Wang, B., Wang, N., Wang, Y., Shen, K., & Li, R. (2022) ‘Characterization of Kinesin Family Member 2C as a Proto-Oncogene in Cervical Cancer’. Frontiers in Pharmacology, 12(January), pp. 1–21. https://doi.org/10.3389/fphar.2021.785981

Yang, L., Zeng, W., Sun, H., Huang, F., Yang, C., Cai, X., Lu, Y., Zeng, J., & Yang, K. (2020) ‘Bioinformatical Analysis of Gene Expression Omnibus Database Associates TAF7/CCNB1, TAF7/CCNA2, and GTF2E2/CDC20 Pathways with Glioblastoma Development and Prognosis’. World Neurosurgery, No. 138, pp. e492–e514. https://doi.org/10.1016/j.wneu.2020.02.159

Yang, Q. X., Zhong, S., He, L., Jia, X. J., Tang, H., Cheng, S. T., Ren, J. H., Yu, H. B., Zhou, L., Zhou, H. Z., Ren, F., Hu, Z. W., Gong, R., Huang, A. L., & Chen, J. (2019) ‘PBK overexpression promotes metastasis of hepatocellular carcinoma via activating ETV4-uPAR signaling pathway’. Cancer Letters, vol. 452(October 2018), pp. 90–102. https://doi.org/10.1016/j.canlet.2019.03.028

Yang, W. X., Pan, Y. Y., & You, C. G. (2019) ‘CDK1, CCNB1, CDC20, BUB1, MAD2L1, MCM3, BUB1B, MCM2, and RFC4 May Be Potential Therapeutic Targets for Hepatocellular Carcinoma Using Integrated Bioinformatic Analysis’. BioMed Research International. https://doi.org/10.1155/2019/1245072

Yin, L., Jiang, L. P., Shen, Q. S., Xiong, Q. X., Zhuo, X., Zhang, L. L., Yu, H. J., Guo, X., Luo, Y., Dong, J., Kong, Q. P., Yang, C. P., & Chen, Y. Bin. (2017) ‘NCAPH plays important roles in human colon cancer’. Cell Death and Disease, vol 8 No. 3, pp. 1–8. https://doi.org/10.1038/cddis.2017.88

Yu, H., Zhang, S., Ibrahim, A. N., Deng, Z., & Wang, M. (2019) ‘Serine/threonine kinase BUB1 promotes proliferation and radio-resistance in glioblastoma’. Pathology Research and Practice, vol. 215 No. 8, pp. 152508. https://doi.org/10.1016/j.prp.2019.152508

Yuan, D., Chen, Y., Li, X., Li, J., Zhao, Y., Shen, J., Du, F., Kaboli, P. J., Li, M., Wu, X., Ji, H., Cho, C. H., Wen, Q., Li, W., Xiao, Z., & Chen, B. (2020) ‘Long non-coding rnas: Potential biomarkers and targets for hepatocellular carcinoma therapy and diagnosis’. International Journal of Biological Sciences, vol. 17 No. 1, pp. 220–235. https://doi.org/10.7150/ijbs.50730

Zhang, B. N., Bueno Venegas, A., Hickson, I. D., & Chu, W. K. (2019) ‘DNA replication stress and its impact on chromosome segregation and tumorigenesis’. Seminars in Cancer Biology, vol. 55(April), pp. 61–69. https://doi.org/10.1016/j.semcancer.2018.04.005

Zhang, H., Yang, X., Zhu, L., Li, Z., Zuo, P., Wang, P., & Feng, J. (2021) ‘ASPM promotes hepatocellular carcinoma progression by activating Wnt / b -catenin signaling through antagonizing autophagy-mediated Dvl2 degradation’. Vol. 11, pp. 2784–2799. https://doi.org/10.1002/2211-5463.13278

Zhang, X., Wang, J., Wang, Y., Liu, G., Li, H., Yu, J., Wu, R., Liang, J., Yu, R., & Liu, X. (2021) ‘MELK Inhibition Effectively Suppresses Growth of Glioblastoma and Cancer Stem-Like Cells by Blocking AKT and FOXM1 Pathways’. Frontiers in Oncology, vol. 10(January), 1–14. https://doi.org/10.3389/fonc.2020.608082

