## Supplementary table for "Identification of prognostic biomarkers for suppressing tumorigenesis and metastasis of Hepatocellular carcinoma through transcriptome analysis"

**Supplementary Table 1** Table showing the detailed analysis of genetic alterations in GBM

| *S. No.* | *Gene Name* | *Types of Genetic Alterations (%)* | *Post Translational Modifications (PTMs)* | *Mutation Type* | *Mutation Site* | *Copy Number Alteration* |
| --- | --- | --- | --- | --- | --- | --- |
| 1 | ASPM | Mutation (2%)  Amplification (6.29%)  Deep Deletion (0.29%) | Phosphorylation  Phosphorylation  Phosphorylation  Phosphorylation, Acetylation, Ubiquitination  Methylation  NA  NA  Phosphorylation, Acetylation, Ubiquitination  NA  Phosphorylation, Acetylation, Ubiquitination  Methylation  Phosphorylation, Acetylation, Ubiquitination  Methylation  Phosphorylation  Phosphorylation, Acetylation, Ubiquitination  Methylation  Phosphorylation,  Acetylation, Ubiquitination  Methylation  Phosphorylation,  Acetylation, Ubiquitination  Methylation  NA  NA  NA | Splice  Nonsense  Nonsense  Nonsense  Missense  Missense  Missense  Missense  Missense  Missense  Missense  Missense  Missense  Missense  Missense  Missense  Missense | L642=  R3152*  R3244*  E2012*  R854H  E918K  S270P  A3137D  T2339P  R2442Q  A3013V  K2586Q  A2152T  D876N  L817F  K2081I  K884M | Diploid  Diploid  Diploid  Diploid  Diploid  Diploid  Gain  Gain  Diploid  Diploid  Diploid  Diploid  Diploid  Diploid  Diploid  Diploid  Diploid |
| 2 | AURKA | Mutation (0.57%)  Amplification (1.43%) | Phosphorylation  Phosphorylation,  Acetylation  Phosphorylation | Missense  Missense  Missense | G11R  R56H  E336D | Diploid  Diploid  Diploid |
| 3 | BUB1 | Mutation (0.57%)  Deep Deletion (0.57%) | Phosphorylation  NA  Phosphorylation,  Ubiquitination  NA  Phosphorylation | Missense  Splice  Missense  Missense  Missense | C698Y  X1021_splice  G533E  F818C  T452M | Diploid  Shallow Deletion  Shallow Deletion  Shallow Deletion Shallow Deletion |
| 4 | BUB1B | Mutation (0.29%)  Deep Deletion (0.57%) | NA | Splice | X950_splice | Diploid |
| 5 | CCNA2 | Mutation (0.29%)  Structural variant (1.43%) | Phosphorylation, Acetylation, Ubiquitination  NA | Missense  Nonsense | A25V  E269* | Gain  Gain |
| 6 | CCNB2 | Amplification (0.57%) | Phosphorylation | Missense | P80S | Shallow Deletion |
| 7 | KIF2C | Amplification (0.29%) | Phosphorylation  Acetylation  Phosphorylation,  Ubiquitination,  Methylation | Missense  Missense  Missense | A648T  E684K  Q508H | Diploid  Diploid  Gain |
| 8 | MELK | Mutation (1.43%) | Phosphorylation | Missense | A315S | Diploid |
| 9 | NCAPG | Mutation (0.86%)  Amplification (0.29%) | Phosphorylation,  Ubiquitination  Phosphorylation  Phosphorylation  NA  Phosphorylation | Missense  Nonsense  Splice  Missense  Missense | A113S  A364L  X373_splice  L444V  S467* | Gain  Diploid  Diploid  Shallow Deletion  Diploid |
| 10 | NCAPH | Mutation (0.29%)  Amplification (0.29%) | NA | Missense | Q704K | Diploid |
| 11 | NUF2 | Mutation (0.79%)  Amplification ( | Phosphorylation,  Ubiquitination  Ubiquitination  Phosphorylation | Missense  Missense Missense | S247Y  Q290H  D435N | Diploid  Diploid  Diploid |
| 12 | PBK | Mutation (0.26%) | Phosphorylation | Missense | Q291K | Diploid |
| 13 | TOP2A | Mutation (0.79%)  Deep Deletion (0.26%) | Phosphorylation, Acetylation,  Sumoylation  Phosphorylation, Acetylation,  Sumoylation  Acetylation, Ubiquitination,  Sumoylation  Phosphorylation, Acetylation,  Methylation,  Sumoylation  Phosphorylation, Acetylation,  Sumoylation | Missense  Missense  Missense  Missense  Missense | T38A  S53Y  V1076F  S1483L  K1492N | Diploid  Diploid  Diploid  Diploid  Diploid |

**Supplementary Table 2** Table showing the detailed analysis of genetic alterations in HCC

| *S. No.* | *Gene Name* | *Types of Genetic Alterations (%)* | *Post Translational Modifications (PTMs)* | *Mutation Type* | *Mutation Site* | *Copy Number Alteration* |
| --- | --- | --- | --- | --- | --- | --- |
| 1 | ASPM | Mutation (2%)  Amplification (6.29%)  Deep Deletion (0.29%) | Phosphorylation  Phosphorylation  Phosphorylation  NA  NA  Acetylation  NA  NA | Missense  Missense  Missense  Missense  Missense  Missense  Missense  FS ins | T111A  K610E  F645L  R792W  I1051T  Y2007S  G2156R  Q2620Tfs*17 | Diploid  Gain  Diploid  Diploid  Gain  Gain  Shallow Deletion  Gain |
| 2 | AURKA | Mutation (0.57%)  Amplification (1.43%) | Phosphorylation  Phosphorylation | Missense  Missense | S83N  R180K | Diploid  Diploid |
| 3 | BUB1 | Mutation (0.57%)  Deep Deletion (0.57%) | NA  NA | Missense  Missense | T805I  S950R | Diploid  Diploid |
| 4 | BUB1B | Mutation (0.29%)  Deep Deletion (0.57%) | NA | Missense | Q460L | Diploid |
| 5 | CCNA2 | Mutation (0.29%)  Structural variant (1.43%) | NA | Missense | N129K | Diploid |
| 6 | CCNB2 | Amplification (0.57%) | NA | No mutation |  |  |
| 7 | KIF2C | Amplification (0.29%) | NA | No mutation |  |  |
| 8 | MELK | Mutation (1.43%) | NA  Phosphorylation  NA  NA | Missense  Missense  Missense  Missense | R53L  I237S/V  V287I  Y638Sfs*4 | Diploid  Diploid  Diploid  Diploid |
| 9 | NCAPG | Mutation (0.86%)  Amplification (0.29%) | NA  NA  Phosphorylation,  Ubiquitination | Missense  Nonsense  Missense | L96F  S574*  Y850H | Diploid  Diploid  Diploid |
| 10 | NCAPH | Mutation (0.29%)  Amplification (0.29%) | NA | Missense | T264A | Diploid |
| 11 | NUF2 | Mutation (0.86%)  Amplification (9.71%) | NA  NA  NA | Missense  Missense Missense | E13D  G385D  Y445C | Gain  Gain  Gain |
| 12 | PBK | Mutation (0.29%)  Deep Deletion (5.71%) | NA | Missense | E303V | Diploid |
| 13 | TOP2A | Mutation (1.14%)  Amplification (0.57%)  Deep Deletion (0.29%) | Phosphorylation  NA  NA  NA | Nonsense  Missense  Missense  Missense | R450*  T689N  R877W  T1315K | Diploid  Gain  Diploid  Diploid |
