## supplementary images for "Identification of prognostic biomarkers for suppressing tumorigenesis and metastasis of Hepatocellular carcinoma through transcriptome analysis"

**Supplementary Figure 1** MA plot representing the log fold-change against mean expression using DESeq2 dataset (a) MA plot of GBM (b) MA plot of HCC

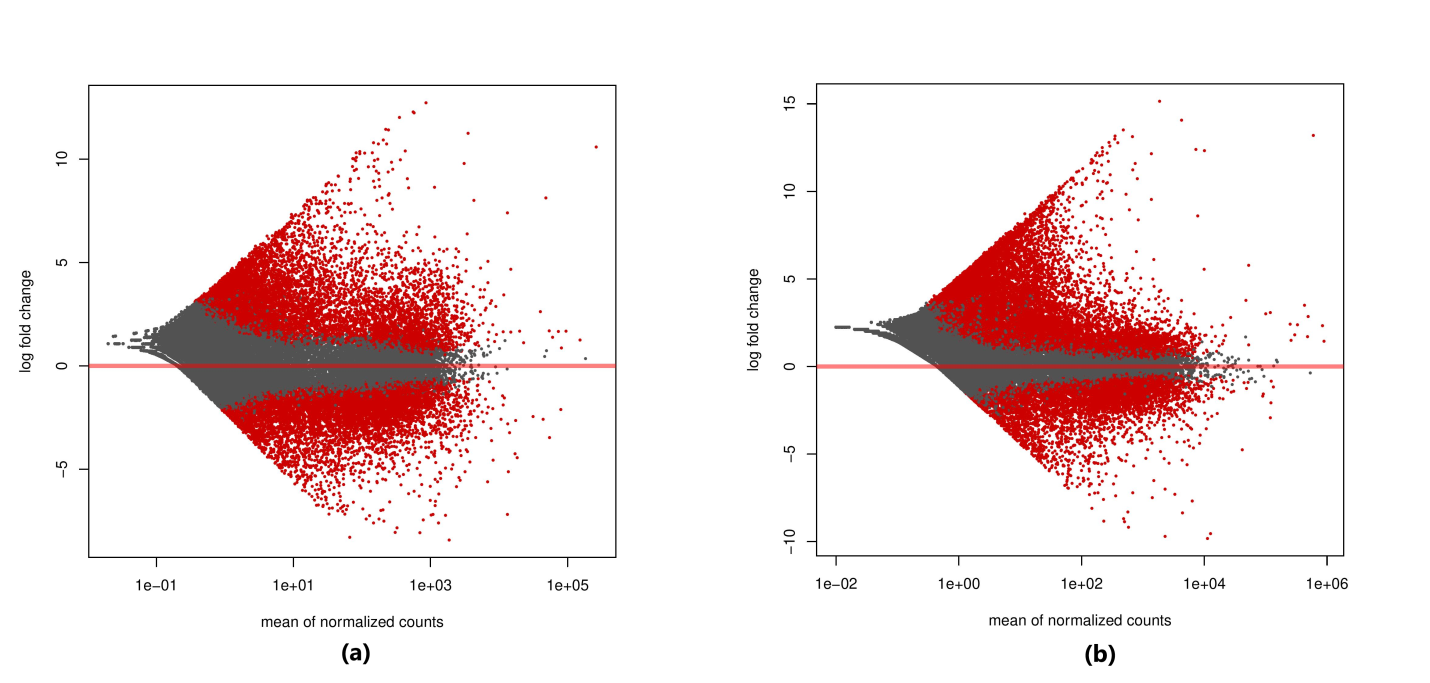

**Supplementary Figure 2** (a) Figure depicting the normal and cancerous brain tissue taken from Human Protein Atlas (HPA) (b) Figure depicting the normal and cancerous liver tissue taken from Human protein Atlas (HPA)

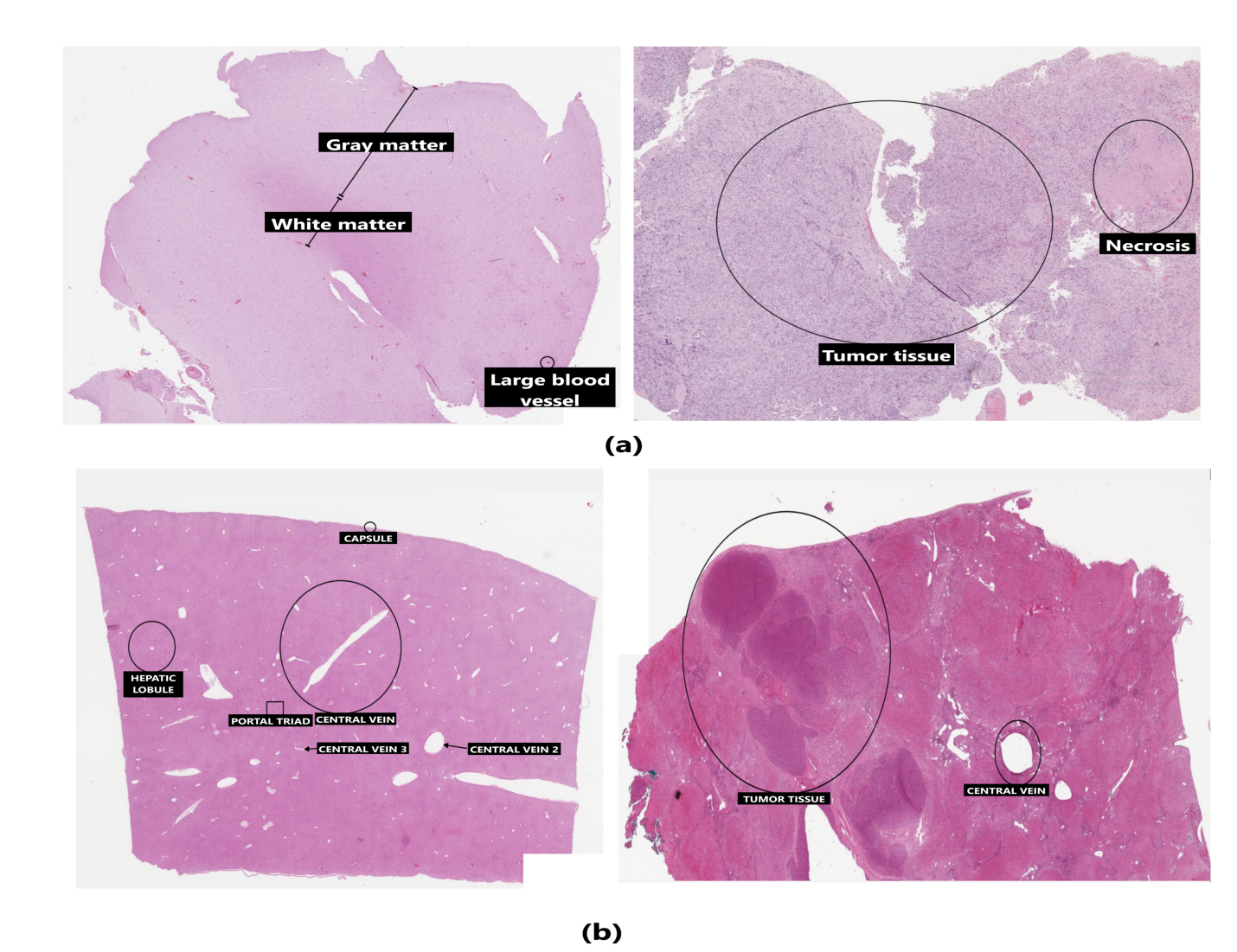

**Supplementary Figure 3** Protein-protein interaction networks of 757 common hub genes between GBM and HCC obtained from DESeq2 analysis. The confidence interval > 0.7. Nodes represent proteins and edges represent interaction between proteins.

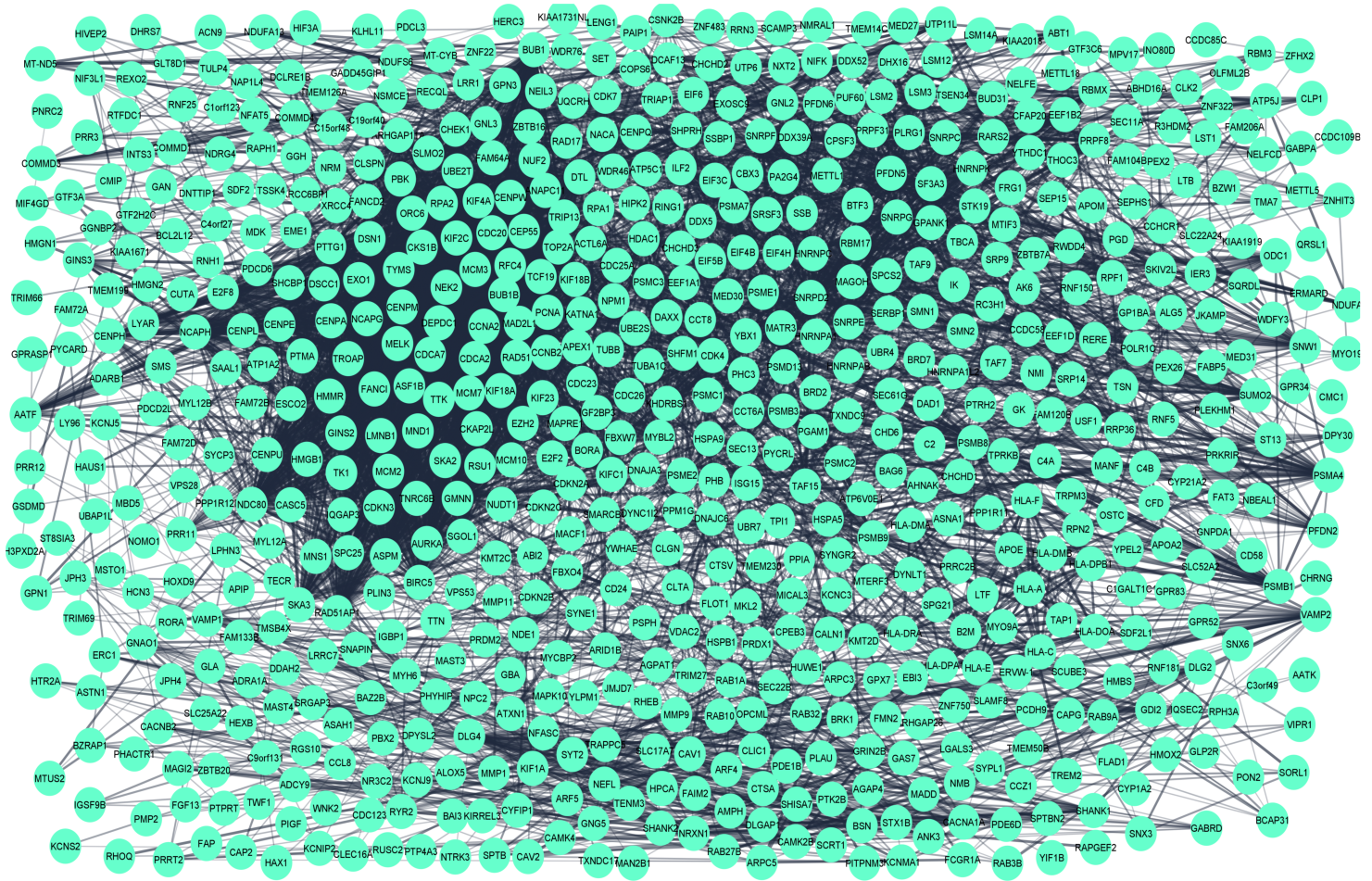

**Supplementary Figure 4 (a)**	Promoter methylation plots of 13 hub genes in GBM

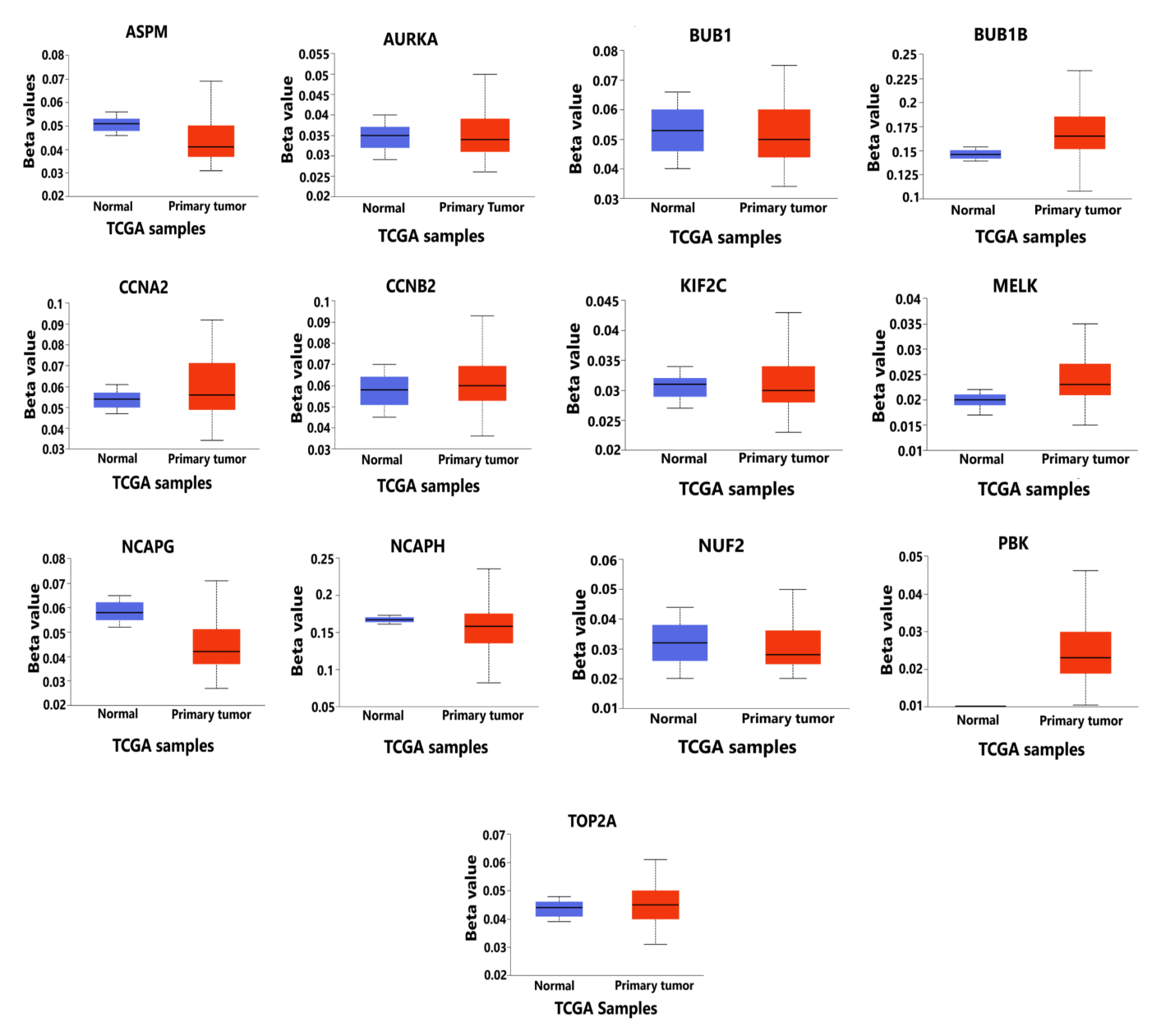

**Supplementary Figure 4 (b)**	Promoter methylation of HCC

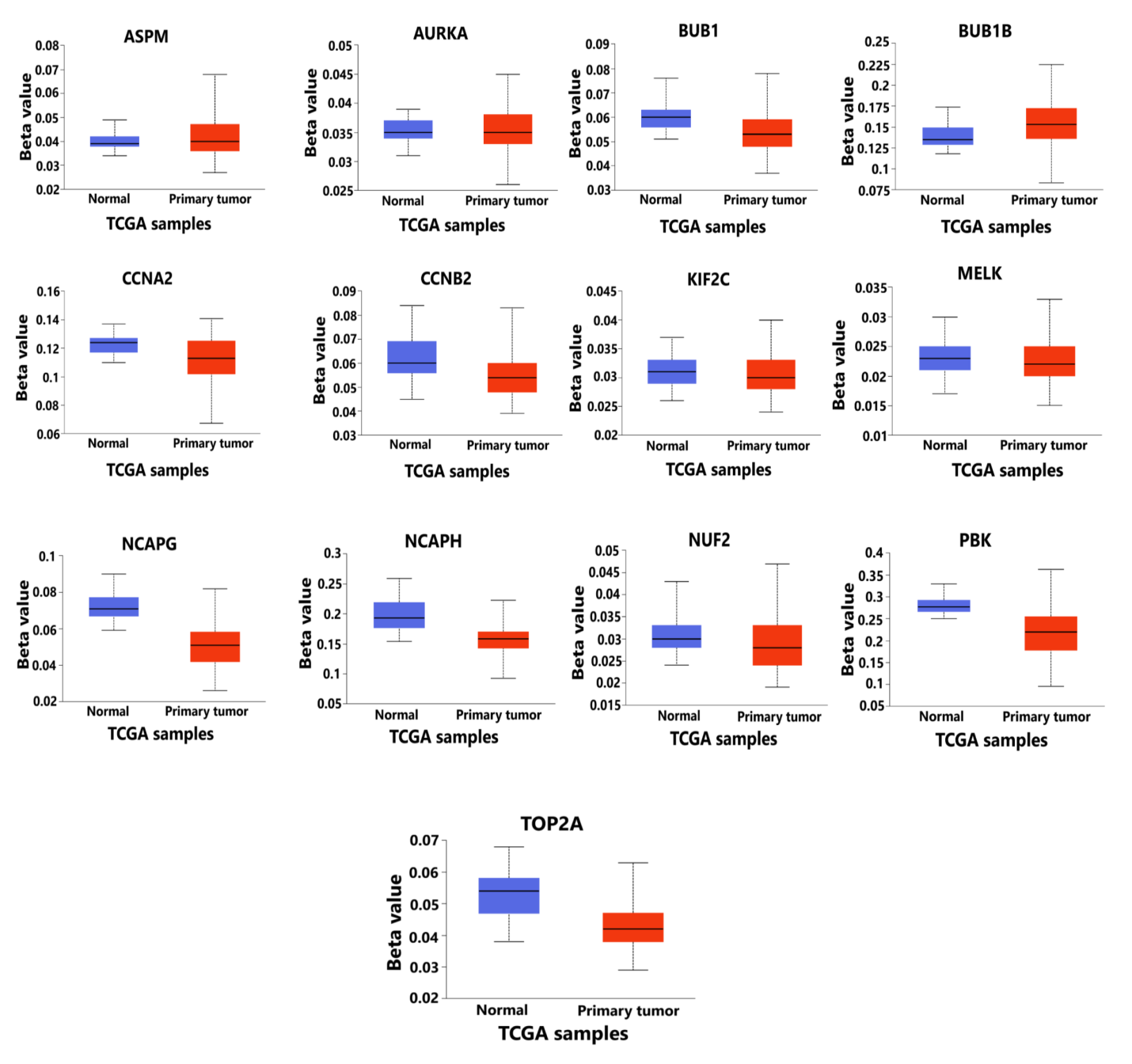

**Supplementary Figure 5** Differential expression pattern of hub genes of both GBM and HCC combined to demonstrate the expression of diseased tumor samples against the normal samples in both the cases. The expression level shows higher expression of samples in case of GBM as compared to those of GBM

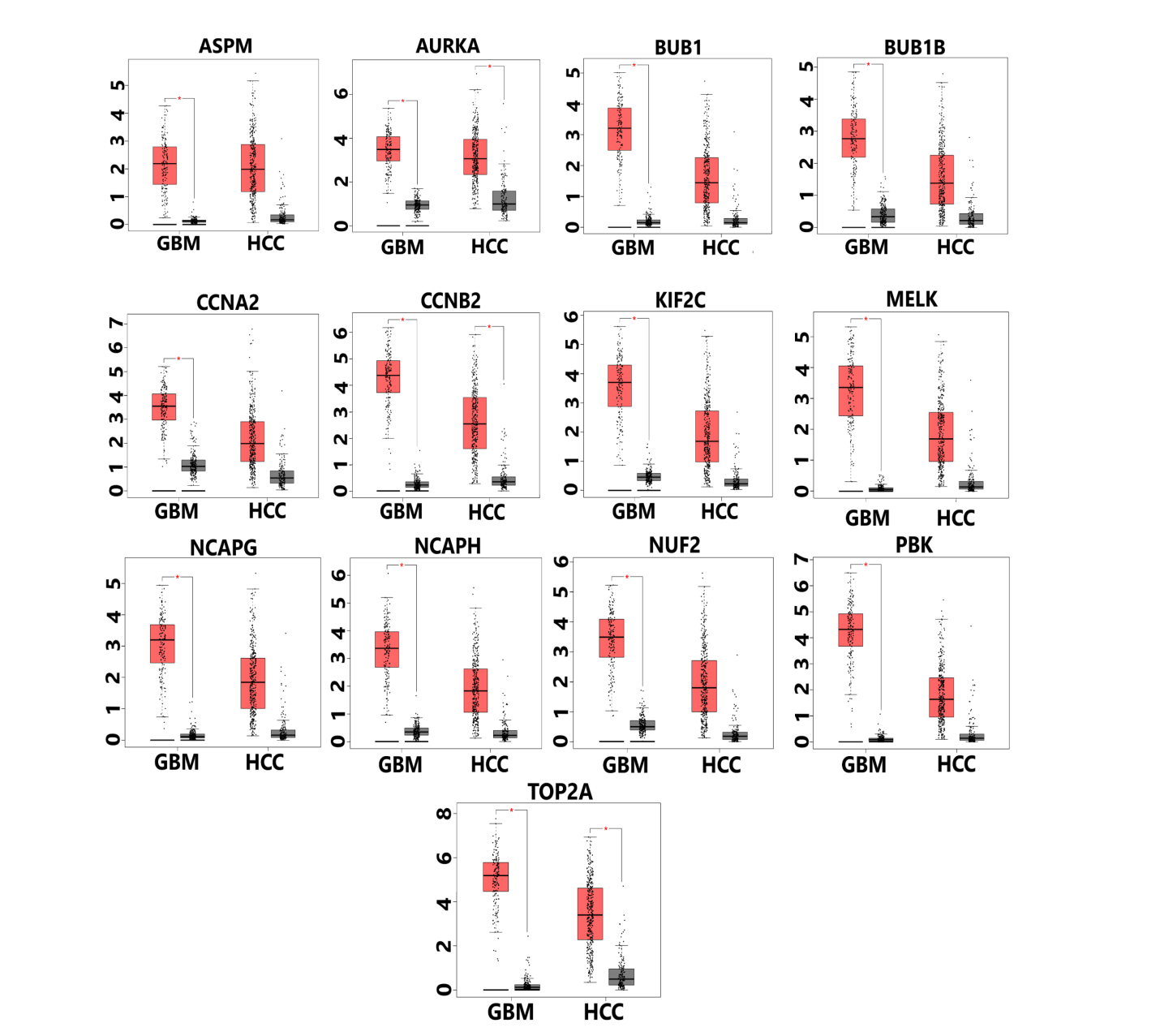

**Supplementary Figure 6** (a) Visualization of genetic alterations of hub genes in GBM using OncoPrint (b) Genetic alterations and their respective alteration frequencies of 13 hub genes in GBM. Green color represents mutations, red color represents amplification, and blue color represents deep deletion. All the hub genes have either missense mutation or nonsense mutation

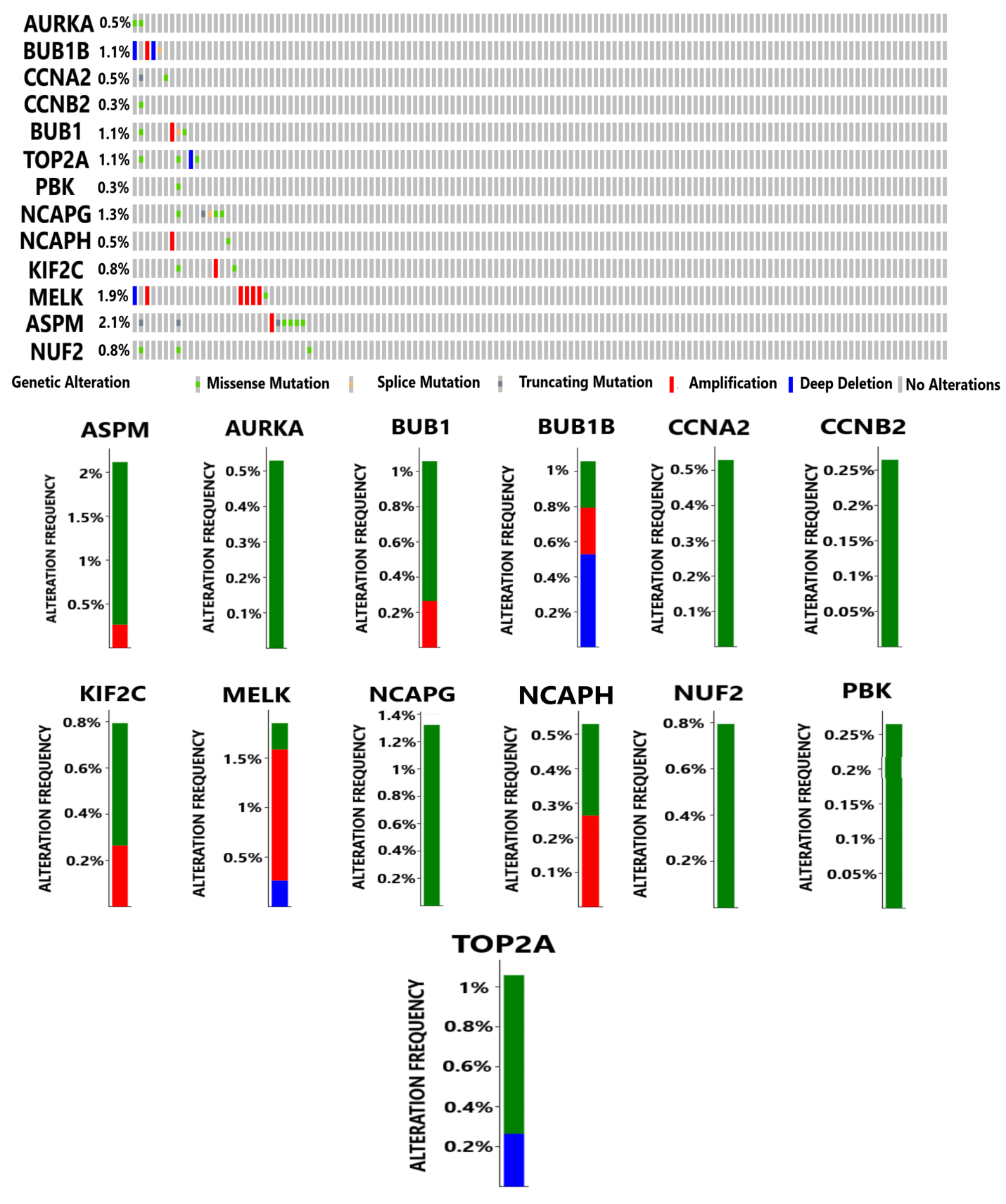

**Supplementary Figure 7**	Copy number variations of 13 hub genes in GBM

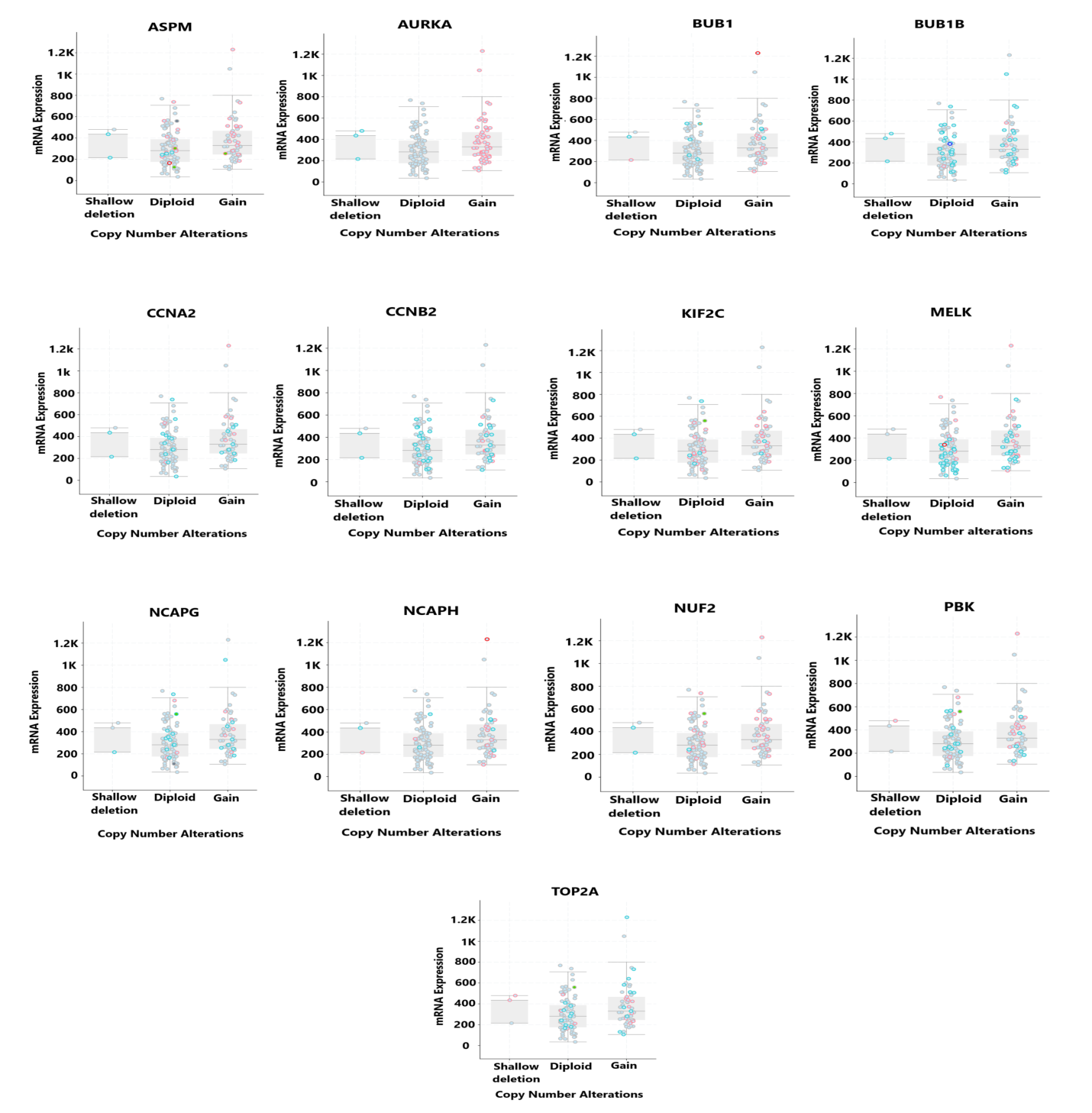

**Supplementary Figure 8** (a) Visualization of genetic alterations of hub genes in HCC using OncoPrint (b) Genetic alterations and their respective alteration frequencies of 13 hub genes in HCC. Green color represents mutations, red color represents amplification, and blue color represents deep deletion. All the hub genes except CCNB2 and KIF2C have either missense mutation or nonsense mutation.

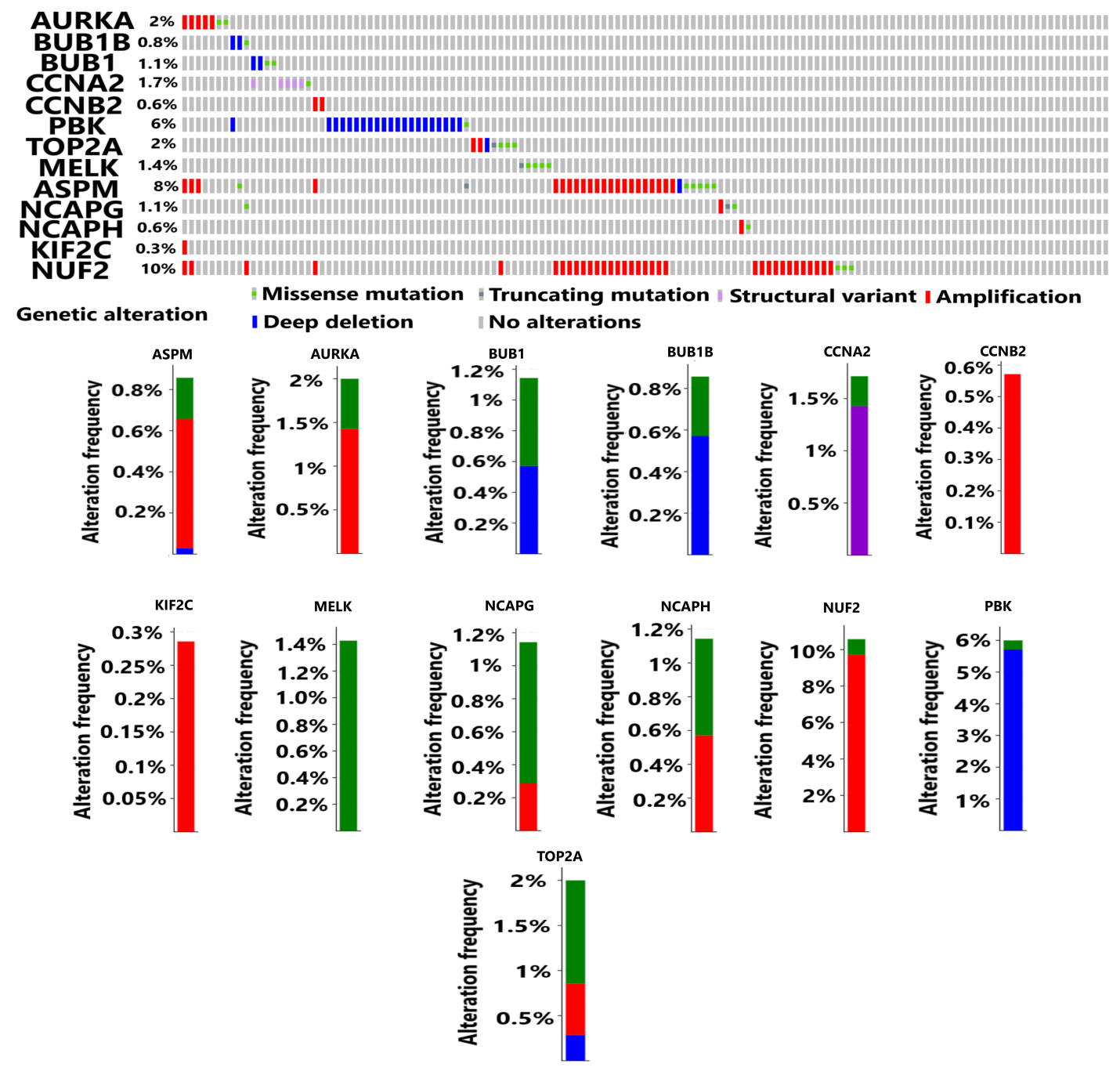

**Supplementary Figure 9**	Copy number alterations of hub genes in HCC

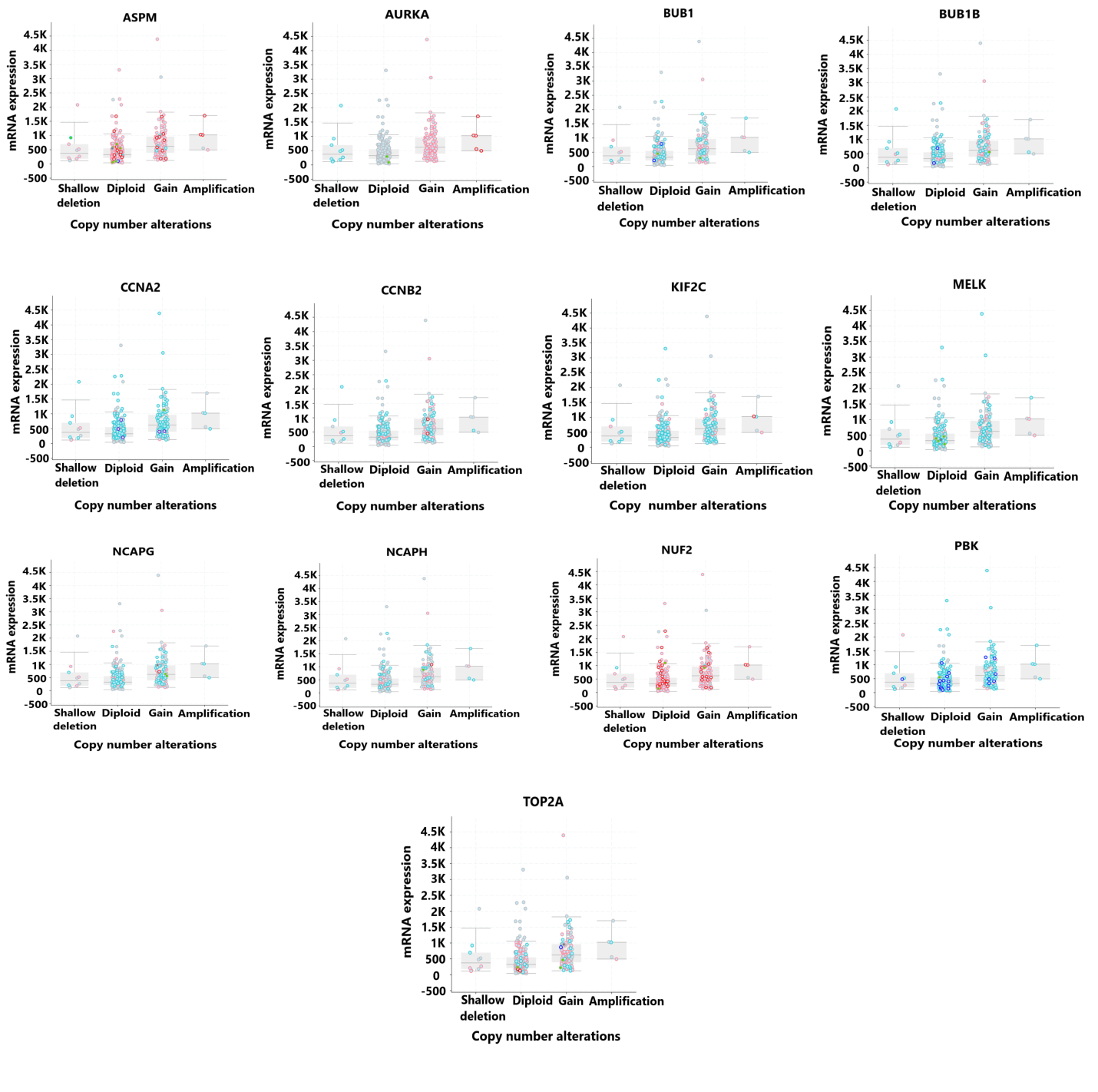
